# Locus Coeruleus Norepinephrine Neurons Facilitate Orbitofrontal Cortex Remapping and Behavioral Flexibility

**DOI:** 10.1101/2023.12.15.571858

**Authors:** M. Cameron Ogg, Hunter T. Franks, Benjamin J. Lansdell, Alex C. Hughes, Jimin Lee, Hunter G. Nolen, Abbas Shirinifard, Lindsay A. Schwarz

## Abstract

To guide behavior, brain regions such as the orbitofrontal cortex (OFC) retain complex information about current tasks and expected outcomes in cellular representations referred to as cognitive maps. When actions produce undesirable results, OFC cognitive maps must update to promote behavioral change. Here, we show that this remapping is driven by the locus coeruleus (LC), a small brainstem nucleus that contains most of the brain’s norepinephrine (NE)-releasing neurons. In a task that tests behavioral flexibility in rodents, LC-NE activity correlated with task acuity and altered depending on trial outcome. Silencing LC neurons caused perseverative behavior and impeded cognitive remapping in OFC, while enhancing LC activity disrupted the ability of new maps to stabilize. These findings reveal a novel role for bidirectional LC-NE signaling in regulation of OFC cognitive map stability and promotion of flexible behavior that differs from the traditional function of this circuit as a global arousal signal.

## INTRODUCTION

To make effective decisions in a changing environment, cellular ensembles in the orbitofrontal cortex (OFC) maintain detailed information about current actions, including available choices, expected outcomes, cues, decision confidence, goals, and outcome value^1–8^. Collectively, the cellular organization of this complex information is often referred to as a cognitive map^9–12^. When behaviors produce unexpected or undesirable consequences, it is important that OFC cognitive maps are updated to reflect the new task contingency so that actions can be altered accordingly. Inhibiting or damaging the OFC produces behavioral deficits in mammalian subjects from rodents to humans that reflect problems in cognitive map updating, such as slowed reversal learning and a failure to adjust behaviors based on the outcomes of their actions^8,11,13^. How OFC cognitive maps are altered to incorporate new information is not fully understood, yet it has been hypothesized to involve the locus coeruleus (LC), a small bilateral brainstem nucleus that contains most of the brain’s norepinephrine (NE)- releasing neurons^14^.

It is theorized that in response to error, or unexpected uncertainty, NE is released in the brain to reset neural networks and shift actions from exploitative to explorative^15–18^. Experimentally, NE has been shown to promote cortical plasticity and behavioral flexibility^19–28^, motivating us to explore how the LC might influence cognitive remapping in the OFC. This is a clinically important topic, as a broad range of neurological conditions that are characterized by cognitive inflexibility (including mood, post-traumatic stress, autism spectrum, attention-deficit and hyperactivity, and obsessive-compulsive disorders, addiction, and Parkinson’s and Alzheimer’s diseases) have also been linked with OFC and/or LC dysfunction^24,29–33^.

To elucidate how LC-NE signaling might influence cognitive remapping and behavior, we established a strategy to simultaneously access the LC and OFC as mice performed a reversal learning task designed to test behavioral flexibility. Fiber photometry of LC-NE neurons revealed temporally precise and outcome-specific activity changes in response to rewards and errors that scaled in magnitude with learning. When LC-NE activity was manipulated during reversal, it disrupted both OFC cognitive remapping (detected via *in vivo* calcium imaging) and behavioral flexibility, though the nature of these deficits was distinct depending on whether LC activity was increased or decreased. While inhibition of LC-NE neurons promoted the retention of prior cognitive maps and perseverative behavior, enhancement of LC-NE signaling impeded the ability of new cognitive maps to settle and task-relevant behavior to emerge. These results define a new role for precise adn bidirectional LC-NE signaling in promotion of OFC cognitive remapping and behavioral flexibility that is distinct from its traditional function as a global arousal signal.

## RESULTS

### LC-NE activity reflects trial outcome and is modulated by learning

To assess LC-NE activity in situations of behavioral flexibility, we established a reversal learning task in which mice learned to navigate an automated T-maze towards one of two lick spouts for sucrose reward delivery (Figure 1A). The calcium indicator GCaMP8m was expressed selectively in LC-NE neurons via injection of a Cre-dependent adeno-associated virus (AAV) into the LC of dopamine beta-hydroxylase-Cre (*Dbh^Cre^*) transgenic mice^34^. A fiber optic cannula was then implanted to chronically monitor bulk calcium-dependent fluorescence changes in the LC as mice navigated the maze during training and reversal sessions across 10 days (Figure 1B, C). During training (T1-T6), mice completed two 6-trial sessions daily and showed improved accuracy at locating the reward spout across training days (Figure 1D). Upon reversal (R1-R4), sucrose delivery was moved to the previously unrewarded spout for subsequent sessions, forcing mice to re-assess the value of each spout and establish a new strategy to obtain reward. Mice initially displayed below-chance performance accuracy early in reversal due to continued navigation to the previously rewarded spout, but performance steadily improved across reversal sessions (Figure 1E).

**Figure 1:**
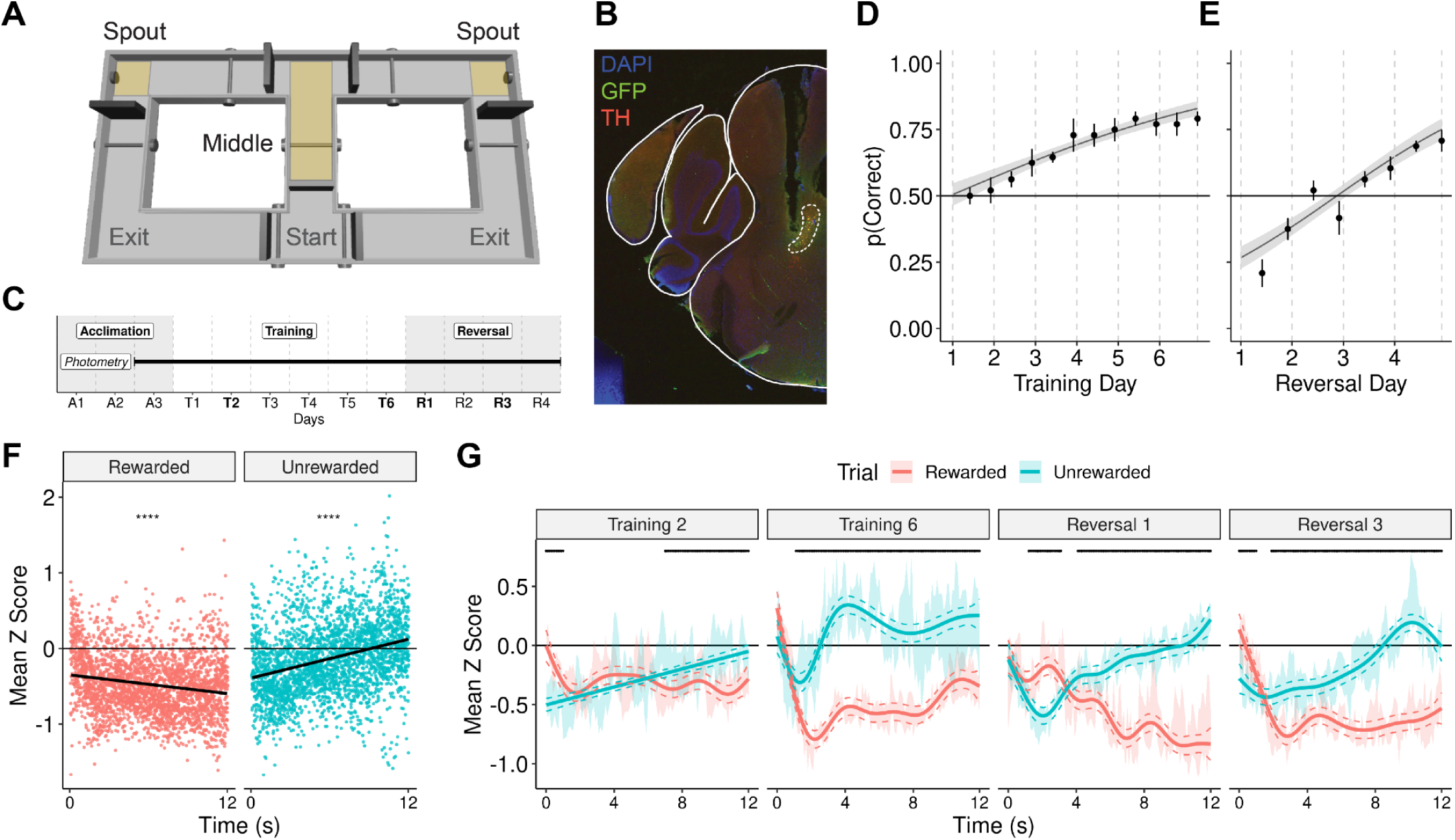
LC activity evolves with behavioral learning and reversal to distinguish trial outcome. (A) Automated T-maze diagram. Regions of interest (spouts/middle) are highlighted in yellow. (B) Coronal section showing the position of the optic fiber and expression of FLEx-jGCaMP8m (green) in TH^+^ (red) LC cells. (C) Experimental timeline schematic. In (D-E), plots show task performance (mean ± s.e.m.): the proportion of rewarded trials/session (dots) and the proportion predicted with a logistic regression model (lines). *n* = 8 mice. (D) Performance during training. Type I ANOVA chi-squared test, trial number: X^2^ (1) = 24.98, p < 0.0001. (E) Performance during reversal. Type 1 ANOVA chi-squared test, trial number: X^2^ (1) = 34.30, p < 0.0001. In (F-G), recorded LC activity is aligned to spout-entry and expressed as the mean *z* score. *n* = 6 mice. (F) Linear regression of LC activity across twelve seconds following spout entry highlights the divergent LC response to trial outcome. Activity decreased at the rewarded spout (r = -0.18) and increased at the unrewarded spout (r = 0.3). The regression significance is displayed on each panel: ****p ≤ 0.0001. Adjusted r^2^ = 0.19 (G) Average LC activity for each trial outcome across days: actual (mean ± s.e.m.; ribbon) and predicted with a generalized additive model (mean ± CrI; lines). Adjusted r^2^ = 0.38. All smooths are significant (p ≤ 0.0001). Significant differences between trial outcomes across time on each day is indicated by the black line.

Analysis of LC-NE activity focused on four days across the reversal learning task: early training (T2), late training (T6), early reversal (R1), and late reversal (R3). Experimental and theoretical studies have shown that, with learning, LC neurons are activated by stimuli predicting reward, rather than the reward itself^18,35,36^. Thus, we thought that the middle of the maze, which the mice navigate prior to altering their trajectory towards a spout, could be a region where anticipatory LC activity develops as mice learn the task structure. Indeed, fiber photometry recordings showed that LC-NE activity increased in this region from T2 to T6, and again from R1 to R3, as the behavior became well-learned and the expectation of receiving a reward was high (Figure S1A, B).

These results suggest that LC-NE neurons signal to predict upcoming rewards, but there is also growing evidence that the LC is activated when these predictions are incorrect^26,37^. It is thought that in response to such unexpected outcomes, as well as to errors in general, release of NE in the brain facilitates neural plasticity and behavioral flexibility^18,28,38^. However, direct measurements of LC-NE activity during explore-exploit tasks are lacking. Fiber photometry paired with our maze task—which is uncued, unforced, self-paced, and freely moving—allows us to directly address this, and on timescales that have previously been unexplored. To investigate how the LC responds to task outcome, we compared LC-NE activity from all animals and all days as they reached the rewarded and unrewarded maze spouts. In line with theoretical predictions, we found that LC-NE activity at the spout changed over time (12 s post-entry) based on trial outcome: increasing following error and decreasing with reward (Figure 1F)^18^. While we observe this general pattern of divergent activity in response to trial outcome across training and reversal, we found that the strength of these responses were modulated by learning (Figure 1G). Early in training (T2), before animals have learned the task, the LC responded similarly to rewarded and unrewarded trial outcomes, with the signals finally diverging 7s after spout entry. This is in stark contrast to later in training (T6), when the unrewarded response was significantly higher than the rewarded response within 1s following spout entry, a difference that was maintained across the time window. A unique pattern of LC-NE activity was also observed on the first day of reversal (R1), where its responses to task outcomes were initially flipped (Figure 1G). Specifically, LC-NE responses at the spout were initially higher for rewarded trials and lower for unrewarded trials (∼1-3s after entry) before ramping up in the following seconds on unrewarded trials only. We believe this reflects a transitory state of LC-NE activity in response to changing task contingencies. As the animals learned the new task structure, LC-NE responses later in reversal (R3) were similar to those seen at the end of training (T6).

In summary, we found that LC-NE activity increased when incorrect choices (errors) were made, and that this response became faster and stronger with learning. Early in reversal (R1), before animals had learned the new task structure, the LC was also briefly excited by unpredicted reward. Throughout the task, when animals made the correct choice, LC-NE activity was selectively suppressed at the outcome of the trial. This effect is unlikely to be driven solely by consumption, as the LC was also briefly suppressed at the unrewarded spout on R1, even though reward was only anticipated^39^. These observations provide new experimental evidence for theories that reductions in LC-NE signaling promote behavioral stability in situations of optimal outcome, while enhanced LC-NE signaling facilitates flexibility when behavior needs to change^14,18^.

### Bidirectional manipulation of LC-NE activity impedes reversal in unique ways

Since we observed that LC-NE activity diverged in response to reward or error and that the magnitude of this difference correlated with learning, we next wanted to examine how manipulating the activity of these neurons altered reversal behavior in the maze task. Pharmacological agents selective for alpha-2 (α2) adrenergic receptors were directly infused into the LC on reversal sessions to transiently silence (clonidine, α2-agonist) or enhance (idazoxan, α2-antagonist) LC activity through an autoreceptor-dependent mechanism (Figure 2A, B)^40,41^. Mice were assigned to one of three treatment groups: clonidine, idazoxan, or control (saline) and received a cannula infusion, according to their group, on each reversal day (R1-R4) before the maze task (Figure 2C). All groups performed similarly during training, when no drugs were administered (Figure 2D). However, pharmacologically manipulating LC activity during reversal sessions significantly reduced performance accuracy, though deficits differed depending on the direction of LC modulation (Figure 2E-G). Inhibiting LC activity induced more severe and longer-lasting preservative behavior (defined as preference for the previously-rewarded spout) in the clonidine group than in controls across the first three days of reversal (Figure 2E, G). In contrast, while performance in control and clonidine-treated animals improved by the end of reversal (R4), performance in the idazoxan group plateaued at chance, with consistently lower accuracy than the control group on R3 and R4 (Figure 2F, G).

**Figure 2:**
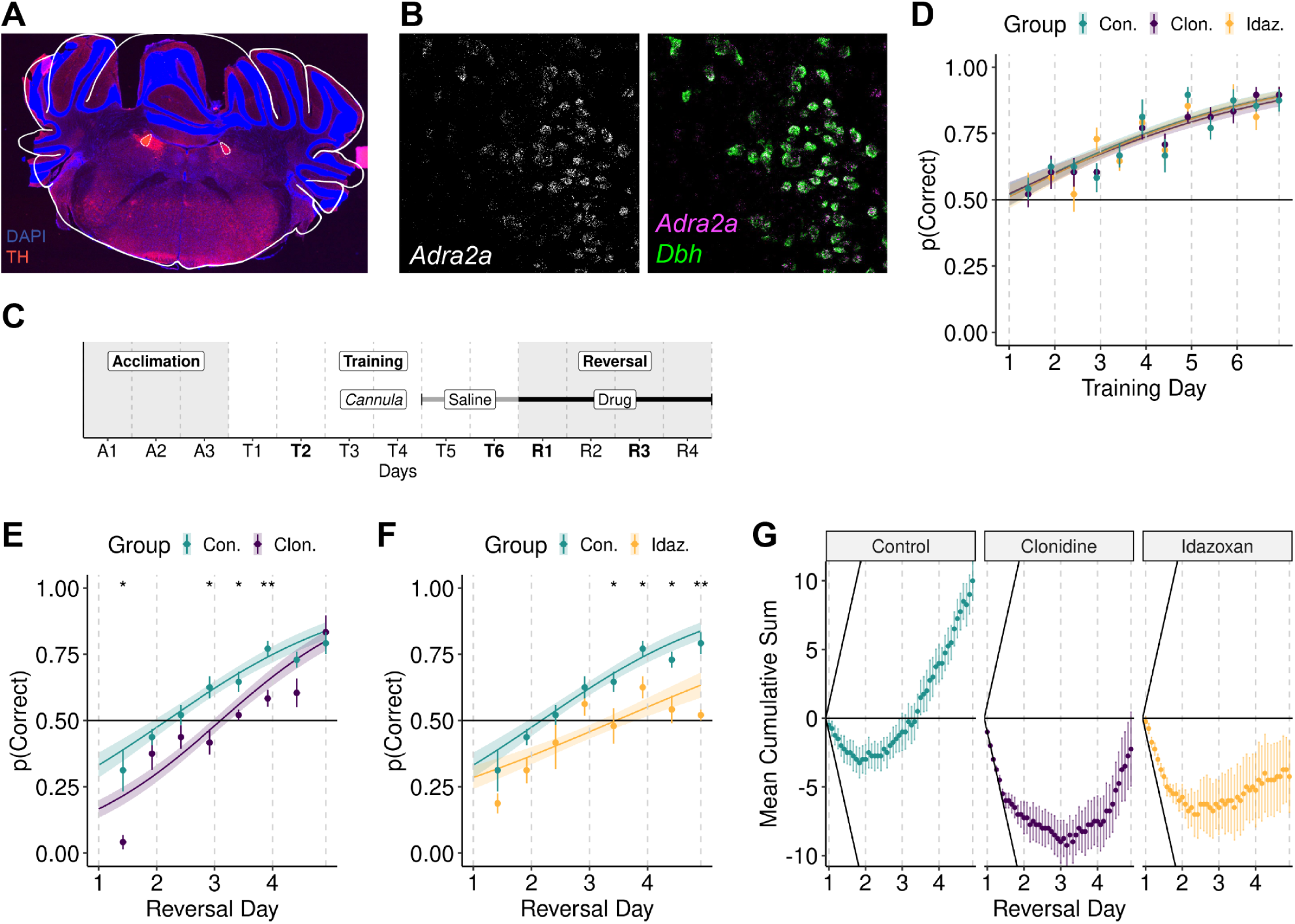
Bidirectional alteration of LC activity impedes reversal learning in distinct ways. (A) Coronal section showing the position of the bilateral cannula above TH^+^ (red) LC cells. (B) *In situ* hybridization of alpha-2A adrenergic receptor (*Adra2a*) transcripts localized to *Dbh*-expressing neurons in the LC. (C) Experimental timeline schematic: Saline was infused on the final days of training. Drug (clonidine, idazoxan, or saline) was infused on each reversal day. In (D-F), plots show task performance (mean ± s.e.m.): the proportion of rewarded trials/session (dots) and the proportion predicted with a logistic regression model (lines). (D) Performance of treatment groups during training. Type 1 ANOVA chi-squared test, trial number: X^2^ (1) = 105.30, p < 0.0001, treatment group: X^2^ (2) = 0.17, p = 0.92, trial number x treatment group interaction: X^2^ (2) = 0.17, p = 0.92. (E, F) Performance of treatment groups during reversal. Type I ANOVA chi-squared test, trial number: X^2^ (1) = 109.87, p < 0.0001, treatment group: X^2^ (2) = 22.04, p < 0.0001, trial number x treatment group interaction: X^2^ (2) = 8.10, p = 0.02. Wilcoxon rank sum test with control as reference, *p ≤ 0.05, **p ≤ 0.01. (E) Comparison of clonidine and control performance. (F) Comparison of idazoxan and control performance. (G) To further visualize behavior, trials were coded based on reward condition (+1 rewarded, -1 unrewarded) and the cumulative sum across all reversal trials was calculated for each animal and averaged across the groups (mean ± s.e.m.). For comparison, diagonal black lines represent the slope of the hypothetical mean cumulative sum if all trials were rewarded (upper) or unrewarded (lower). *n* = 7 control, 8 clonidine, 8 idazoxan mice.

In addition to cannula pharmacology, we also manipulated LC-NE neurons using chemogenetics. Excitatory (3D) or inhibitory (4D) DREADDs (Designer Receptors Exclusively Activated by Designer Drugs) were expressed in LC-NE neurons via injection of a recombinase-dependent AAV into the LC of *Dbh^Cre^* or *Dbh^Flp^* transgenic mice (Figure S2A). Post-experimental histology revealed varying DREADD expression levels in these animals, despite maintenance of consistent viral injection conditions across all experiments (Figure S2B). To activate DREADD-expressing LC-NE neurons, mice were given an intraperitoneal injection of the agonist clozapine N-oxide (CNO) before reversal sessions (Figure S2C). All groups performed similarly during training, when no CNO was administered (Figure S2D). During reversal, results in the chemogenetic groups were similar, but not as robust, as those in the cannula pharmacology groups (Figure S2E, F). We suspect this difference might be due to variability of DREADD expression within the LC, which could reduce the robustness or reliability with which LC-NE activity was modulated in these animals (Figure S2C).

Notably, comparing reversal performances of DREADD and pharmacologically-manipulated animals revealed an inverted U-shaped effect of NE signaling on behavior, similar to that defined in the Yerkes-Dodson Law, whereby too little or too much arousal impedes optimal performance^23,42^. One explanation for this phenomenon is that disruption of NE signaling—in either direction—affects the stability of behaviorally-relevant cellular ensembles in downstream brain areas such as OFC^14,18^. Indeed, we confirmed that LC-NE neurons synapse directly onto excitatory (*Emx1^+^*) cells within OFC, positioning them to directly modulate OFC neural activity Figure S3A-D). Based on these results, we next wanted to ask how bidirectional changes in LC-NE activity might influence cognitive map structure in OFC.

### OFC forms cellular representations of behavioral state that remodel when task contingencies change

Before assessing how LC-NE signaling influences OFC cognitive map stability, we first needed to characterize the formation, spatial organization, and stability of task-relevant cell ensembles within this region throughout the course of the maze task. Cre-dependent GCaMP7s-expressing AAV was injected into the OFC of *Emx1^Cre^* transgenic mice, followed by two implants: a cannula into the LC for drug infusion (as described previously) and a GRIN lens into the OFC (Figure 3A). Using a microendoscope, we chronically recorded activity from thousands of GCaMP-expressing excitatory OFC neurons (mean = 340 cells per mouse, range = 158-713 cells per mouse) as mice underwent training and reversal sessions (Figure 3B). Tracking cells over multiple days (Figure S4A-C), we observed an ∼70% overlap of responsive cells from day to day, with individual cells active on a variable number of those days (Figure S4D). All cells were included in subsequent analyses, regardless of how many days they responded.

**Figure 3:**
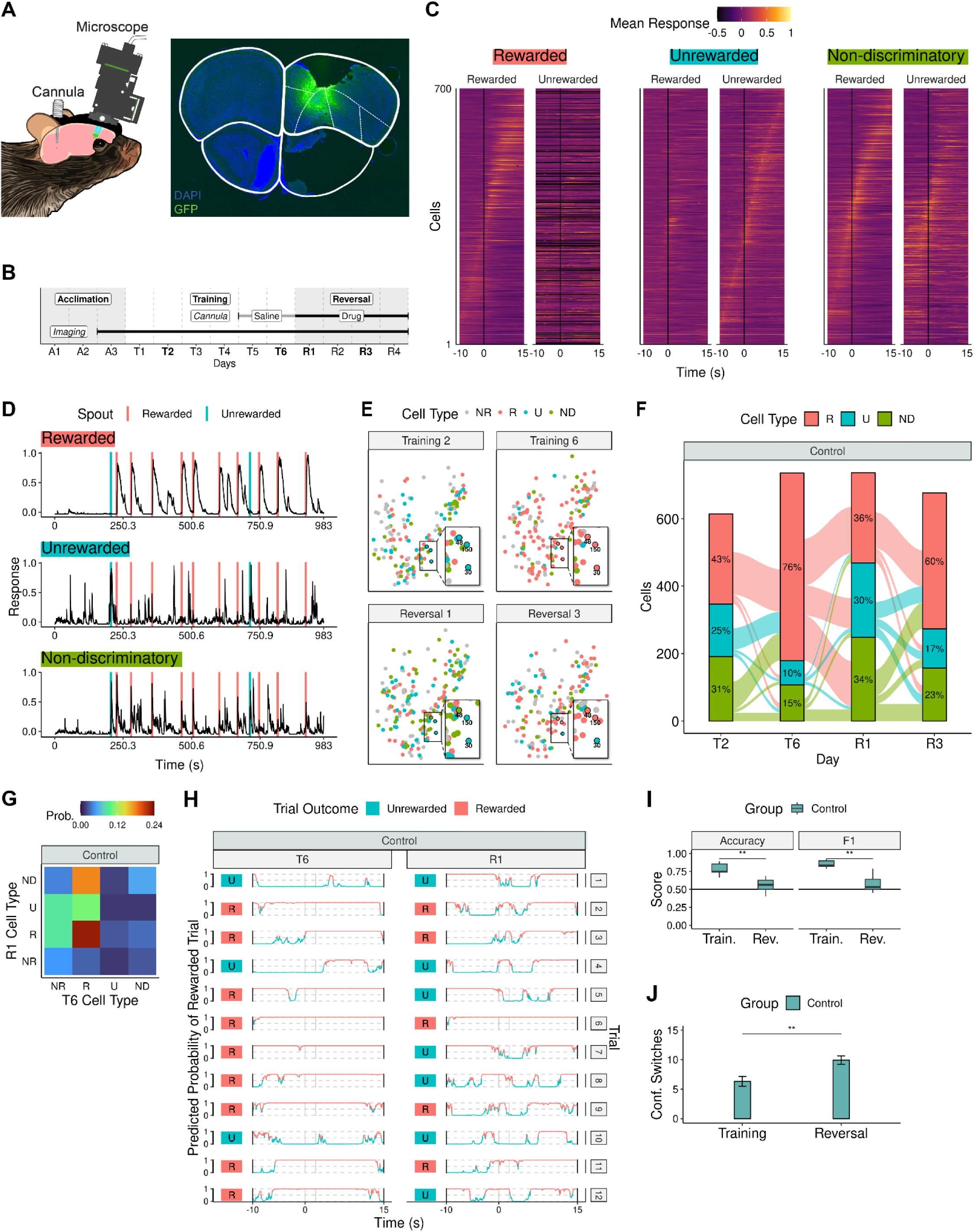
OFC task representation during training and reversal. (A) Left, diagram of LC cannula and OFC lens implants. Right, coronal section showing the position of the GRIN lens and expression of FLEx-GCaMP7s (green) in the OFC. (B) Experimental timeline schematic. (C) Trial-averaged responses during training of 700 OFC cells of each ROC-identified cell type randomly selected from all animals (sorted by time of maximal response). *n* = 9 animals. (D) The entire T6 response of one OFC cell of each type from an example control animal. Vertical colored lines mark initial spout entry for each trial. (E) Spatial location of OFC cells from an example control animal colored by cell type (NR = Non-responsive). Inset highlights how cell type changes across the task in three cells that present on all four days. (F) Alluvial plot showing how the distribution of cell types changes during training and reversal in the control group. *n* = 3 mice. (G) Joint probability distribution between T6 cell type and R1 cell type for an example control animal with significant mutual information. (H) Trial-by-trial decoder prediction across time on T6 and R1 for an example control animal. The colored box next to each trace indicates the actual trial outcome (rewarded/unrewarded). Vertical gray lines represent the 2-second period in which the decoders’ predictions were tested. (I) Metrics comparing the quality of the control decoders in predicting trial outcomes during training and reversal. Box limits, first and third quartiles; center line, median; whisker limits, minimum and maximum values; points, data exceeding box limits ± 1.5x interquartile range. Decoder performance was impaired during reversal. Wilcoxon signed rank test, **p ≤ 0.01. (J) Average number of confidence switches/trial in the control decoder predictions across time (mean ± s.e.m.). Confidence switches increased from training to reversal. Wilcoxon rank sum test, **p ≤ 0.01.

The OFC cognitive map is thought to include cellular ensembles that predict and evaluate behavioral outcomes^13^. In our task, the middle of the maze represents a pre-choice region where predictions about task outcome might occur and the spouts are post-choice regions where outcomes can be evaluated. Fiber photometry recordings showed that LC-NE activity changed in these regions of the maze (Figure 1F, G; Figure S1A, B), motivating us to focus on these regions for further analysis. To identify OFC cellular ensembles that are responsive within these regions, receiver operating characteristic curve (ROC) analysis was used to identify cells with selective activity in the middle of the maze (trial initiation) and/or at either spout (trial completion) (Figure S5A-D). Across all animals and days, 73% of OFC cellular recordings contained task-related responses, meaning that a cell’s activity changed significantly in these regions of interest compared to its activity in the rest of the maze. OFC cells were classified into three major types based on their ROC-identified responses: *Rewarded*, *Unrewarded*, or *Non-discriminatory* (*ND*) (Figure 3C, D). *Rewarded* cells significantly increased or decreased activity only during rewarded trials. The opposite was observed for *Unrewarded* cells, where significant changes in activity (increased or decreased) occurred during unrewarded trials. *ND* cells had significant activity changes in the same direction during both rewarded and unrewarded trials.

While ROC-identified OFC cells responded with high reliability within a given day, we wondered how these ensembles might evolve with learning or when outcomes are suddenly altered. Thus, we next looked at how these cellular ensembles change throughout the maze task. The proportion of *Rewarded* cells grew across training, recruiting both new cells and those that had previously been in the *Unrewarded* or *ND* ensembles, thereby strengthening the OFC representation of the rewarded side of the maze (Figure 3E, F). On R1, there was also an increase in the proportion of *Unrewarded* cells that selectively responded to the previously rewarded side of the maze, reflecting continued encoding of the previous task structure within OFC (Figure 3E, F). However, many cells also began transitioning their responses on R1, either becoming *ND* cells, potentially reflecting an intermediate map state as cells progress towards reward-encoding, or *Rewarded* cells (Fig. 3e, f). In support of this, when we calculated the probabilities that T6 cells of a certain cell type become R1 cells of a certain cell type (i.e. a joint probability distribution), the highest probabilities were that a T6 *Rewarded* cell would become a R1 *Rewarded* or a R1 *ND* cell (Figure 3G). As reversal progressed and the proportion of *Rewarded* cells increased (Figure 3E, F), the OFC cognitive map continued to remodel to reflect the new task state. Using mutual information (MI) to quantify how much information about one state (e.g. reversal cognitive map) can be determined from another state (e.g. training cognitive map), we found that the number of control animals that had significant MI between their training and reversal maps decreased from R1 (2/3) to R3 (1/3)(Methods).

Since cognitive maps are theorized to guide behavior, we next tested whether task outcome could be accurately decoded from the collective activity of these cellular ensembles. We trained a linear decoder on two seconds of OFC population activity at spout entry during training trials (T2 and T6). When tested on T6, individual decoders could determine whether a training trial was rewarded with high accuracy and F1 score (a measure of precision and recall), indicating that the OFC cognitive map forms an accurate task representation (Figure 3H, I). However, when the task changed in reversal sessions, the accuracy and F1 score of these same decoders significantly decreased, implying that the previous (T6) cognitive map no longer accurately encodes the reward status of a trial (Figure 3I). When we applied the decoders across a larger time window surrounding spout entry for each trial (–10s to +15s), we observed that predictions regarding whether a trial was rewarded (pink) or not (blue) fluctuated between periods of high and low confidence (>99% or <1%) (Figure 3H). We speculate that these confidence switches may reflect consideration of available choices, as previously reported,^1^ as well as uncertainty in outcome evaluation. In support of this, we observed that when behavior was well-learned and the accuracy of the decoders was high, these switches were less frequent. When the task changed with reversal, the number of confidence switches in our decoders’ predictions significantly increased (Figure 3H, J). Collectively, this is evidence that the previous OFC cognitive map fails to appropriately guide behavior when task contingencies change.

### Perturbation of LC-NE signaling alters OFC cognitive map updating upon reversal

To evaluate the effect of LC-NE activity on the organization of OFC cellular ensembles during reversal, we compared the distribution of cell types among our treatment groups to those seen in a control reversal (Figure 4A, B). Inhibition of the LC during reversal resulted in a higher proportion of *Unrewarded* OFC cells in clonidine animals on both reversal days (Figure 4A, B; Figure S6A, B), in line with their perseverative navigation to the previously rewarded side of the maze (Figure 2E, G). Further, nearly half of reversal (R1 and R3) *Unrewarded* cells in these mice had been *Rewarded* cells on T6 (Figure S6C), showing that with reduced LC-NE activity, OFC cellular ensembles from training remain active as reversal proceeds. This differs from what was seen in animals where LC-NE activity was increased (via idazoxan) during reversal, where the majority of reversal (R1 and R3) *Unrewarded* cells were not part of the previous *Rewarded* cell ensemble (Figure S6C, D). Additionally, as reversal progressed (R3) and performance declined, the idazoxan group had a higher proportion of *ND* cells (Figure 4B, Figure S6E, F). These results indicate that chronic enhancement of LC-NE activity increases the responsiveness of OFC cells to both task-relevant and irrelevant information, potentially making it harder for an accurate OFC cognitive map and behavioral strategy to form and stabilize. As further evidence that poor performance in the clonidine group was driven by continued activity of the T6 cognitive map, a joint probability distribution revealed that a T6 *Rewarded* cell had the highest probability of becoming a R1 *Unrewarded* cell (Figure 4C). Additionally, all clonidine animals had significant MI between training (T6) and both reversal days, indicating that with loss of LC-NE signaling, the OFC cognitive map does not remodel appropriately. In contrast, MI between training and reversal was found in only a single idazoxan animal on a single day (R1), with the highest joint probability occurring between T6 *Rewarded* and R1 *ND* cells (Figure 4C). Thus, with enhanced LC-NE signaling, the T6 OFC cognitive map is less persistent throughout reversal.

**Figure 4:**
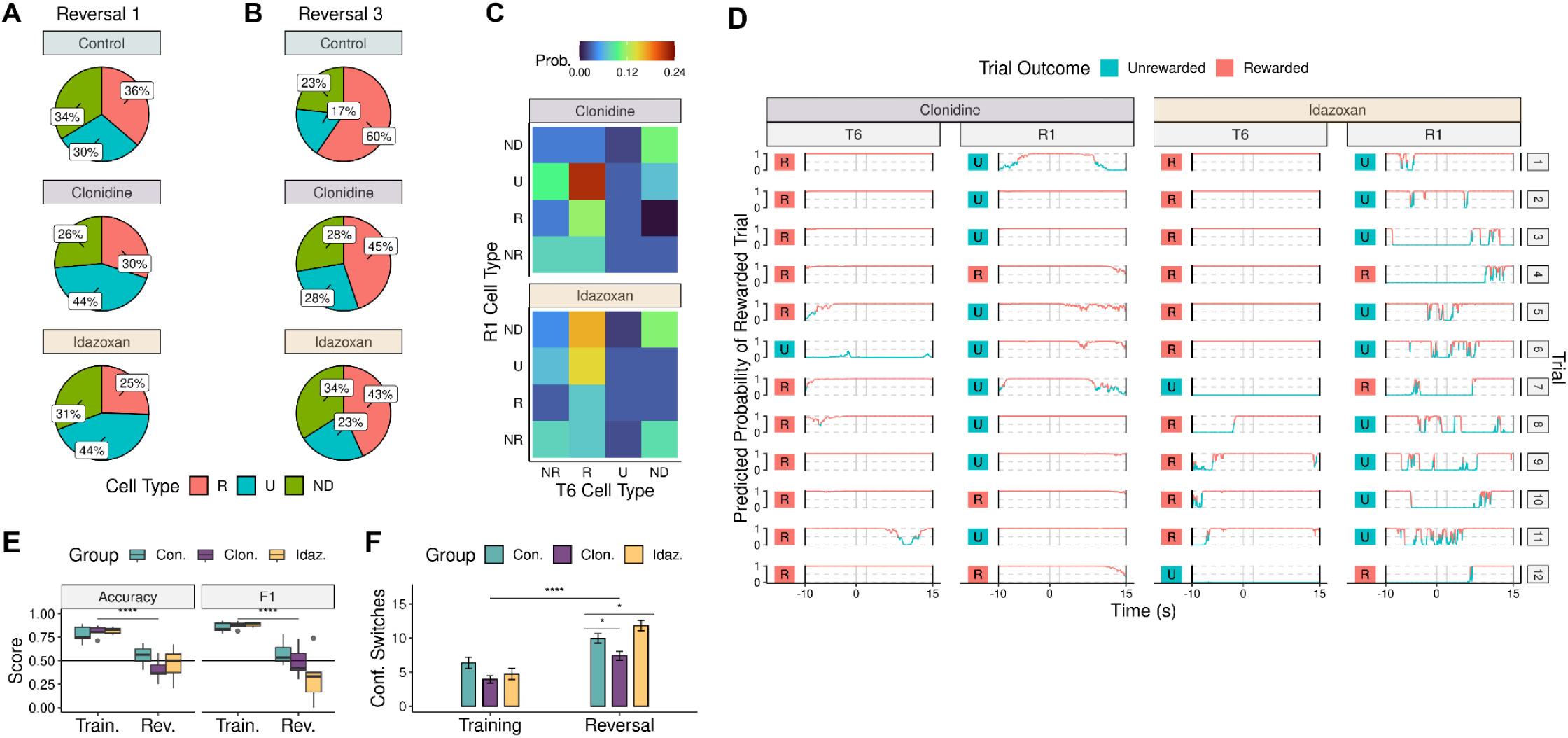
OFC task representation is affected by bidirectional alteration of LC activity. In (A-B), plots show the distribution of cell types across treatment groups during reversal. (A) R1 comparison. Pearson’s chi-squared test, X^2^ (4) = 44.10, p < 0.0001. (B) R3 comparison. Pearson’s chi-squared test, X^2^ (4) = 56.19, p < 0.0001. (C) Joint probability distribution between T6 cell type and R1 cell type for an example animal with significant mutual information in each treatment group. **(D)** Trial-by-trial decoder prediction across time for an example animal in each treatment group on T6 and R1. The colored box next to each trace indicates the actual trial outcome (rewarded/unrewarded). Vertical gray lines represent the 2-second period in which the decoders’ predictions were tested. (E) Metrics comparing the quality of the decoders in predicting trial outcomes during training and reversal. Box limits, first and third quartiles; center line, median; whisker limits, minimum and maximum values; points, data exceeding box limits ± 1.5x interquartile range. Decoder performance was impaired during reversal. Wilcoxon signed rank test, ****p < 0.0001. These metrics did not differ between treatment groups. Kruskal-Wallis rank sum test, Training Accuracy: p = 0.69, Reversal Accuracy: p = 0.38, Training F1: p = 0.56, Reversal F1: p = 0.10. **(F)** Average number of confidence switches/trial in the decoder predictions across time (mean ± s.e.m.). Confidence switches increased from training to reversal. Wilcoxon rank sum test, ****p < 0.0001. Confidence switches did not differ among the groups during training. Kruskal-Wallis rank sum test, p = 0.10. Confidence switches differed among the groups during reversal. Kruskal-Wallis rank sum test, p < 0.0001. Wilcoxon rank sum test with control as reference, Clonidine: *p = 0.02, Idazoxan: *p = 0.04. *n* = 3 control, 3 clonidine, 3 idazoxan mice.

Finally, we examined how LC-NE signaling alters OFC cognitive map representation of task state during reversal by assessing the ability of decoders to determine trial outcome in clonidine and idazoxan groups. As in control animals, decoders trained on OFC population activity during training trials performed worse and had more confidence switches in their predictions during reversal (Figure 4D-F). However, we observed differences in OFC decoder predictions between treatment groups that again emphasize distinct origins for the behavioral deficits of these animals, now at a network level. The clonidine decoders had fewer confidence switches across the 25 second window of analysis (Figure 4D, F) and a higher rate of false positives (Figure S6G) than the control group, supporting evidence from the ROC and MI analyses that reduced LC-NE activity impairs OFC cognitive map updating. This suggests that clonidine animals evaluate the outcome of unrewarded trials incorrectly and may consider the newly rewarded side of the maze as a behavioral choice less often. In contrast, decoders in the idazoxan group predicted that 50% of trials were rewarded, regardless of the animals’ actual behavior (Figure S6H). Idazoxan decoders also had more confidence switches than those of the control group (Figure 4F), implying that increased LC activity puts the OFC in a plastic state of alternating between predictions, rather than settling into a new task state.

## DISCUSSION

Many decades of research indicate that OFC plays a critical role in guiding adaptive behavior across mammalian species, with recent theories proposing that this process involves maintenance of an accurate cognitive map within OFC^10,43,44^. Advances in large-scale monitoring of neural activity *in vivo* are facilitating discovery of the cellular components underlying these abstract representations^4,5,45,46^, but less is known about how the brain updates these maps to maintain optimal behavioral output when the environment changes. Neurons within OFC can switch their response properties within several trials of a task contingency change, suggesting that they receive cues almost immediately after an unexpected outcome to begin encoding new information (Figure 3F)^47^. It seems unlikely that such a signal originates solely from within the OFC, as these neurons do not robustly encode prediction errors^48–50^. Instead, neuromodulatory circuits, especially dopamine and serotonin, have been implicated in driving behavioral flexibility, though conflicting data exists regarding their specific influence within OFC^51–59^. Here, we provide experimental evidence that a different neuromodulatory circuit, specifically NE-releasing neurons in the LC, activate in response to unexpected or undesirable outcomes to facilitate the remapping of neural ensembles within OFC and promote flexible behavior.

While, to our knowledge, these are the first experiments to directly implicate LC-NE neurons in OFC cognitive remapping, these findings harmonize with theoretical predictions that LC might play such a role^14,18^. Using fiber photometry, we observed that LC-NE activity distinguishes task outcome, responding robustly to errors and unexpected outcomes and suppressing activity in the presence, and even the expectation, of reward (Figure 1F, G). Recent computational work on LC function—which modeled LC activity similar to what we experimentally observed in our recordings—posited that when behavior needs to change, increased LC activity facilitates cognitive map flexibility, improving task performance and enabling reversal learning^18^. Indeed, when we inhibited normal LC-NE dynamics using either chemogenetic (4D) or pharmacological (clonidine) manipulation, reversal learning was negatively impacted (Figure 2E, G; Figure S2E, F). We observed that this behavioral deficit correlated with a greater proportion of OFC neurons responding to the previously rewarded side of the maze, causing the cognitive map to inaccurately represent the current task structure (Figure 4, Figure S3, Figure S6).

Our observation that pharmacologically increasing LC activity (idazoxan) also negatively impacted reversal learning (Figure 2F, G; Figure S2E, F) might be viewed as incongruent with studies showing that enhancement of NE signaling can heighten attention, facilitate attentional shifts, and increase brain-wide functional connectivity^20,21,60,61^. Intriguingly, several animals that underwent chemogenetic activation of LC-NE neurons (3D) had the best reversal performances that we observed from any of the animals, including controls (Figure S2F; light green dots). However, there was also particularly high variability in the reversal performance of individual 3D-expressing animals. We speculate that these effects could be due to individual differences in the level of DREADD expression within the LC of these mice, which in turn produced variability in the level to which LC-NE activity was enhanced (Figure S2B). Thus, we think that the behavioral differences seen within manipulations (Figure 2F, G; Figure S2E, F), and between our study and others, reflect an inverted U-curve effect, where intermediate levels of LC-NE activity promote optimal behavioral output^15,23^. Ultimately, the precise level of NE signaling that is needed to optimize cognitive function is unknown. However, experiments where LC activity was enhanced during reversal (Figure 2F, G; Figure S2E, F), as well as our fiber photometry results (Figure 1F, G), suggest that a key parameter for accurately modulating cognitive maps in OFC might be the timing of LC activity. When LC activity is sustained independently of task outcome, as occurred in our idazoxan group, a higher proportion of OFC neurons become responsive to both task-relevant and irrelevant information (Figure 4, Figure S6). This leads to a cognitive map that cannot settle into a new task state, promoting continued exploratory behavior.

Understanding the mechanisms by which NE-related circuits maintain such a nuanced balance of activity is critical when considering that deficits within these circuits are implicated in many neurological diseases. Recent studies have demonstrated that enhancement of LC activity can improve performance in animals with cognitive^24,25^ or auditory deficits^62^, emphasizing broad clinical relevance for NE-targeted therapeutics. Yet, for other diseases where NE-related brain circuits are strongly implicated, such as anxiety and depression, treatments that alter NE signaling have shown limited efficacy^63,64^. Our findings here, that both the level and timing of NE signaling influence cortical mapping and cognitive performance, suggest that these features, which likely differ across diseases and possibly even across patients—are important to consider when leveraging the LC-NE system as a therapeutic target.

## Data availability

The datasets used for the figures can be obtained from the authors.

## Code availability

Code can be obtained from the authors.

## Acknowledgements

We thank Maïlys Ayerdi and Hollie Sanders for technical support; Robbie Hughes for developing custom maze control software and designing the maze graphic; Alec Ogg for designing the mouse graphic for Fig. 3; Pat Stemkowski for Inscopix imaging support; and Stanislav Zakharenko, Jan Lui, Max Fletcher, and Schwarz lab members for helpful feedback on the manuscript. This work was supported by institutional funds from St. Jude Children’s Research Hospital (M.C.O., H.T.F., B.J.L., J.L., H.G.N., A.S., L.A.S.), and funding from the St. Jude Graduate School of Biomedical Sciences (A.C.H).

## Author Contributions

M.C.O. and L.A.S. designed the project. L.A.S. generated viruses, performed in situ hybridization and rabies tracing experiments, and supervised the project. M.C.O. performed the chemogenetic, pharmacology, and OFC calcium imaging experiments, with assistance from H.T.F. and H.G.N. on behavior and histology. A.C.H., H.T.F., and J.L. performed the LC fiber photometry experiments. A.S. advised on imaging analysis and supported data management. M.C.O. and B.J.L. performed data analysis. M.C.O. and L.A.S. wrote and edited the manuscript with feedback from the other authors.

## Methods

### Animals

Adult (>3 mo) male and female mice (C57BL/6, *Emx1^Cre^*, *Dbh^Cre^* KI, and *Dbh^Flp^*) were group-housed up to 5 mice per cage, maintained on a 12:12-hr light/dark cycle (lights on at 06:00) with ad libitum food and water. Experimental protocols were approved by and conformed to St. Jude Children’s Research Hospital IACUC and conform to US National Institutes of Health guidelines.

### Behavior

An automated T-maze (Fig. 1A, 55cm, 105cm, 20cm (L,W,H), Maze Engineers) task was used to assess reversal learning behavior. Mice were given access to water ad libitum but were put on a restricted diet starting 2-3 days before the task began, which reduced and maintained body weight at ∼85-90% of its original value throughout the duration of the experiment. Mice received food following task completion each day. Prior to behavioral testing, mice were habituated to any task components that involved handling.

The task, adapted from^65^, included three phases (Fig. 1c): acclimation, training, and reversal. Each day (except for A1), the mouse was put into the start area and allowed to habituate for 2-5 minutes before starting the task. The maze was cleaned between mice with 70% ethanol.

#### Acclimation

On A1, mice were acclimated to sucrose and drinking from the spouts. 40% sucrose was loaded into spouts on both sides of the maze and small drops were also placed on the floor around the spouts. All maze doors were closed and the mouse was put in one of the arms with a spout. The mouse stayed in that arm until it had consumed sucrose from the spout, at which point it was moved to the alternate maze arm. On A2/A3, mice were acclimated to the automated doors and the task structure. 40% sucrose was loaded into spouts on both sides and the mouse completed 6 trials (1 session) on each day.

#### Training

On T1-T6, 40% sucrose was loaded into only one spout and the mouse completed 12 trials (2 sessions, separated by a 1 min break) on each day. To be included, mice had to average at least 75% correct trials across the final two training days.

#### Reversal

On R1-R4, 40% sucrose was loaded into the opposite spout from that used during training and the mouse completed 12 trials (2 sessions, separated by a 1 min break) on each day.

During the task, the position of the mice in the maze was recorded using AnyMaze video tracking software (Stoelting Co.). Afterwards, the position of the mice was verified manually. For analysis, we defined several regions of interest (ROI): middle (vertical stem after start and before arms, 35×10 cm) and spout (area in front of spout at the end of each arm, 10×10 cm) (Fig. 1a).

### Stereotaxic surgeries and virus injections

For all stereotaxic surgeries, mice were anesthetized with isoflurane (2.5-3%, 0.75 L/min), and received an analgesic dose of Marcaine (0.5 mg/ml, 0.1 ml s.c. at scalp, St. Jude Pharmacy), and were placed on a heating pad and in a stereotaxic apparatus (David Kopf Instruments, 922) where anesthesia was maintained with Somnoflo nose-cone isoflurane (2%, 0.2 L/min, Kent Scientific, SF-01). Following surgery and for the next 3 days, mice were given Meloxicam (0.5 mg/ml, 1-2 mg/kg s.c., St. Jude Pharmacy) for pain relief. Mice were given at least 3 weeks to recover after surgery before starting experimental protocols.

Virus injections were made using a microinjection pump (World Precision Instruments; MICRO-2T) and pulled glass electrodes (Drummond Scientific Company, 5-000-2005). 400 nl of virus was injected at a rate of 40 nl/min, unless otherwise specified. The electrode was not removed until 5 minutes after the end of injection to allow diffusion of the virus.

### Viral tracing

For rabies tracing experiments, AAV8-hSyn-FLEx^loxP^-TVA-mCherry and AAV8-hSyn-FLEx^loxP^-H2B-P2A-N2cG-WPRE were mixed at a 1:1 ratio prior to injecting 200 nl into the left OFC of *Emx1^Cre^* (The Jackson Laboratory, 005628) mice. 2-3 weeks later, 200 nl of RABV-CVS-N2c^ΔG^-H2B-eGFP (titer: ∼2.1E+8 infectious particles per ml) was injected into the same location within OFC. One week later, mice were perfused and brains were collected for histology of putative starter cells within the OFC and GFP-expressing neurons within the LC.

### *In situ* hybridization

Mice were euthanized with Avertin (12.5 mg/ml, 240 mg/kg i.p., St. Jude Pharmacy) and fresh brain tissue was harvested and immediately mounted in OCT for cryosectioning. 16-20 µm tissue sections containing the LC were collected for RNAscope-based fluorescence in situ hybridization (ACDBio). Samples were labeled using the RNAscope Multiplex Fluorescent Reagent Kit v2 (ACDBio), and following the supplied protocol. The following probes were used: Adra2a-C1 (425341) and Dbh-O1-C3 (464621-C3). Probe signal was developed using Opal dyes (Opal 520 and 570 Reagent Pack, Perkin Elmer) at a dilution of 1:1000 and counterstained with DAPI. Fluorescent images were taken using a Leica LSM780 laser scanning confocal microscope.

### Immunohistology

To verify viral targeting and implant placement, mice were euthanized with Avertin (12.5 mg/ml, 240 mg/kg i.p., St. Jude Pharmacy) and transcardially perfused with 1x PBS (10 ml) and 4% PFA (10 ml) after completion of experiments. When confirming implant placement, the head was kept overnight in 4% PFA. Brains were removed, fixed overnight in 4% PFA, cryoprotected overnight in 30% sucrose, mounted in OCT, and cut on a cryostat (Leica CM1950) in 40 μm slices.

Antibody staining was performed on tissue sections attached to SuperFrost Plus microscope slides (Fisherbrand). Slides were washed in 1x PBS (10 min) and then covered with a blocking buffer (1x PBS with 0.3% Triton-100 (PBST) and 10% NDS) and Parafilm (30 min). Slides were incubated overnight in primary antibody (diluted in PBST and 5% NDS) at room temperature and then washed 3x (10 min) in PBST. Slides were incubated for 2 hours in secondary antibody (diluted in PBST and 5% NDS) at room temperature and then washed 2x (10 min) in PBST, 1x (10 min) in PBS, and 1x (10 min) in PBS and DAPI. Glass coverslips were mounted to the slides with Fluoromount-G (SouthernBiotech). All images were captured at 5x magnification with a Leica DFC 3000G microscope.

Primary antibodies used were: rabbit anti-Tyrosine Hydroxylase (TH) (ab152, Millipore, 1:1000), mouse anti-TH (mab318, Millipore, 1:500), chicken anti-GFP (GFP-1020, Aves Lab, 1:2000), and rabbit anti-RFP (600-401-379, Rockland, 1:1000). Secondary antibodies used were: Cy3 anti-rabbit (711-165-152, Jackson, 1:1000), Cy3 anti-mouse (715-166-151, Fisher, 1:500), 488 anti-mouse (NC0192065, Jackson, 1:500), and 488 anti-chicken (703-545-155, Jackson, 1:1000).

For histology examples in the paper, outlines were created by warping a matched section from the Paxinos and Franklin atlas (Franklin and Paxinos 2019; Gaidica) to the example section using the BigWarp ImageJ plugin^66^. LC outlines were determined by TH-antibody staining.

### Fiber photometry

To record LC activity during the task, *Dbh^Cre^* KI (The Jackson Laboratory, 033951) mice (n=4) were injected with a Cre-dependent virus encoding GCaMP8 ( AAV-Syn-FLEx-jGCaMP8m, titer: 2.11*10^13; St. Jude Viral Vector Core) into the left LC (AP (from lambda): 1.0, ML: -0.8, DV (from brain surface): 3.15 and and implanted with a unilateral fiber optic cannula (ferrule size: 1.25mm, core diameter: 200μm, length: 4mm, NA: 0.37, Neurophotometrics) above the virus injection (DV (from brain surface): 3.05).

During the task, excitation illumination (wavelengths: 465 nm, 405 nm) was generated though fiber-coupled LEDs and modulated by a real-time amplifier (Tucker-Davis Technologies, RZ10x). Excitation lights were filtered and combined by a fluorescence mini cube (Doric, FMC6). The combined excitation light was delivered through a mono fiber optic patch cord (core diameter: 200μm, NA: 0.37, Doric, D207-1310-2) to the implanted fiber optic cannula. Fluorescence was sampled at 1000 Hz. Synapse software (Tucker-Davis Technologies) was used to record the fiber photometry data and control the LEDs. The optimal LED power for each animal, not exceeding 60 mA, was determined before experiments began based on a Synapse quality measurement.

Traces were exported from Synapse and aligned to the behavioral videos in Matlab. In R, the traces (465 nm signal) were downsampled to 10 Hz, the ΔF/F was calculated using the isosbestic traces (405 nm signal) as baseline to correct for bleaching and artifactual signal fluctuations as in^67^ and converted to z scores as in^39^. For each animal and day, we extracted parts of the trace before (8 s) and after (12 s) entry into the middle or the spout and averaged these matched segments across trials of the same type (rewarded or unrewarded). Statistical analysis was performed in R. For the behavioral data, we fit two logistic regression models (glm, family=binomial) to the training and reversal behavioral data. We used a deviance goodness-of-fit test (ANOVA Chi-square) to assess whether trial significantly affected the fit of the models. To assess where across time the LC response to trial outcome on each day was significantly different, we fit a generalized additive model (gam, family=scat) to the spout-aligned data and compared the difference between predicted smooths using simultaneous confidence intervals, calculated based on 10000 simulations.

### Cannula pharmacology

To assess how changing the activity of the LC during reversal learning affects performance, C57BL/6 (The Jackson Laboratory; 000664) mice (n=22) were singly-housed following stereotaxic surgery to implant bilateral guide cannulae (26 GA, 2.8 mm length, 2 mm separation, P1 Technologies, 8IC235GS520S) above the LCs (AP (from lambda): -1.0, ML: ±1.0 ,DV (at skull): -2.8). Custom injectors (33 GA, P1 Technologies, 8IC235IS5SPC) projected 1 mm past guide cannulae.

On days of infusion (Fig. 2B), 600 nl of 0.9% sodium chloride (Hospira, St. Jude Pharmacy), α2-antagonist idazoxan hydrochloride (5 mg/ml, adapted from^68^, Sigma, I6138), or α2-agonist clonidine hydrochloride (5 mM, adapted from^40^, Sigma, C7897) were infused bilaterally (200 nl/min, 2 min wait) using a dual-syringe (5 μl, Hamilton, 87931) microinjection pump (KD Scientific, 14-831-211) and plastic tubing (P1 Technologies, 8F023X041P01) immediately before the maze task. Testing prior to experiments determined drug dosages that did not cause gross impairments in behavior.

Statistical analysis was performed in R. We fit two logistic regression models (glm, family=binomial) to the training and reversal behavioral data. We used a deviance goodness-of-fit test (ANOVA Chi-square) to assess whether trial or NE treatment group significantly affected the fit of the models. For each session, performance (mean percent correct trials) was compared using a Wilcoxon test with the control group as reference.

### Chemogenetics

To further assess how changing the activity of the LC during reversal learning affects performance, *Dbh^Cre^* KI mice or *Dbh^Flp^* mice (The Jackson Laboratory; 033952)(n=40), were injected (350 nl) with a Cre- or Flp-dependent virus encoding either excitatory or inhibitory Designer Receptors Exclusively Activated by Designer Drugs (DREADDs) (pAAV-hSyn-DIO-hM3D(Gq)-mCherry (AAV5), titer: 1.3 × 10^13, AddGene, 44361); pAAV-hSyn-DIO-hM4D(Gi)-mCherry (AAV5), titer: 1.2 × 10^12, AddGene, 44362); hSyn-FLEx-FRT-3D-mCherry (AAV8), St. Jude Viral Vector Core; hSyn-FLEx-FRT-4D-mCherry (AAV8), St. Jude Viral Vector Core) bilaterally into the LC (AP (from lambda): 1.0, ML: -0.8, DV (from brain surface): 3.15. DREADD expression was verified post-histology, and any animals without notable expression were included in a Miss group.

For acclimation to the injection, mice were administered 0.9% sodium chloride (0.1 ml i.p.,Hospira, St. Jude Pharmacy) 30 mins prior to the task at the end of training (T5, T6). For DREADD activation on reversal days, mice were administered clozapine N-oxide (CNO) (1 mg/ml, 5 mg/kg i.p., Hello Bio, HB6149) 30 mins prior to the task.

### Calcium imaging

To record OFC activity during the task, *Emx1^Cre^* mice (n=9) were singly-housed following stereotaxic surgery to inject (30 nl/min) a Cre-dependent virus encoding GCaMP7 (DJ-CAG-FLEx-GCaMP7s, Schwarz lab) into the right OFC (AP: 2.5, ML: 1.1, DV (at skull): -1.8) and to implant a Proview GRIN lens (diameter: 1 mm, length: 4 mm, Inscopix, 1050-004605) over the virus injection (DV (at skull): -1.6). The skull was removed using a 1 mm-diameter biopsy punch (info) and a small amount of brain tissue was aspirated above the implantation area. Once in place, the lens was surrounded by Kwik-Sil (World Precision Instruments, 50822154) to protect the surrounding brain tissue and attached using super glue and dental cement. These mice were also implanted with bilateral LC cannulae, as described above. After around 3 weeks, a baseplate (Inscopix) was installed above the lens with super glue and dental cement at a height where responsive cells were in focus.

During the task, imaging data was recorded at 20 Hz using a microendoscope (Inscopix, nVista 3.0), Inscopix Data Acquisition Software, and the AnyMaze digital interface (Stoelting Co.). Videos were resized in Inscopix Data Processing Software and exported to CaImAn software^69^ for motion correction. CaImAn was used to identify responsive cells, match cell identity across days, and export imaging traces. Imaging traces were normalized as the ratio (F-F_median_)/(F_max_-F_min_) as in^4^ and aligned to the behavioral videos in R.

#### ROC-AUC

To analyze single cell responses during the maze task, responses (*z*-scored calcium traces) of single neurons in each maze ROI (rewarded/unrewarded spout, rewarded/unrewarded middle) were quantified using ROC (receiver operating characteristic) analysis as in^70^, using custom R scripts. For each neuron, a threshold is applied to its neural activity, producing a binary vector (1 if above threshold, 0 if below). A binary vector denoting behavior is also generated (1 if the animal was in the ROI, and 0 if not). These vectors are compared to determine a true positive rate (TPR) and a false positive rate (FPR) over all time-points. Plotting the TPR against the FPR over a range of thresholds, spanning the minimum and maximum values of the neural signal, yields an ROC curve that describes how well the neural signal detects behavior events at each threshold. We used the area under the ROC curve (auROC) as a metric for how strongly neurons were modulated in our ROIs. For each neuron/ROI, the observed auROC was compared to a null distribution of 1,000 auROC values generated from constructing ROC curves over randomly permuted calcium signals (traces that were circularly permuted using a random time shift). A neuron was considered significantly inhibited in a given ROI if its auROC value was less than the 5^th^ percentile of the random distribution and significantly excited if it exceeded the 5^th^ percentile.

Each cell was assigned a cell type on each day it was recorded based on its significant responses: *Non-responsive* cells did not have significant activity changes in the ROIs during either trial type. *Rewarded* cells had significantly increased or decreased activity during rewarded trials. *Unrewarded* cells had significantly increased or decreased activity during unrewarded trials. *Non-discriminatory* cells had significant activity changes in the same direction (increased/decreased) during both rewarded and unrewarded trials. A small subset of cells had significant activity changes in the different directions for rewarded and unrewarded trials and were assigned as *Rewarded* or *Unrewarded* based on their increased response (e.g. a cell that had increased activity on rewarded trials and decreased activity on unrewarded trials was assigned to the *Rewarded* group) .

On each day, cells from all animals in each treatment group were compiled and the overall proportions of cell types were compared across groups using a Pearson’s chi-squared test. Individual proportions were compared between and across groups using a pairwise proportion test adjusted for multiple comparisons.

#### Mutual Information

To analyze the relationship between a cell’s activity (i.e. cell type) during training and reversal, we performed a mutual information (MI) analysis^71^ using custom Python scripts. For a given mouse, cells that could be reliably identified between T6, R1 and R3 were used. The mutual information between the cell type on T6 and cell type on R1 was computed. The same was done for T6 and R3. A resampling procedure was used to determine what mutual information values were significantly above chance. Labels were shuffled 100 times to generate a null distribution of mutual information values. An alpha value of 0.05 was used to determine what was a significant amount of mutual information. The number of mice with significant mutual information between T6 and reversal are summarized in the table:

| Group | Reversal Day | % (n/3) |
| --- | --- | --- |
| Control | 1 | 67 (2) |
| Control | 3 | 33 (1) |
| Clonidine | 1 | 100 (3) |
| Clonidine | 3 | 100 (3) |
| Idazoxan | 1 | 33 (1) |
| Idazoxan | 3 | 0 (0) |

#### Decoder

To measure population-level encoding of reward status by OFC neurons, we used custom Python scripts to construct linear discriminant analysis (LDA) decoders to predict the reward status (rewarded vs. unrewarded) of trials based on population activity. Training sets and test sets were constructed using population vectors evoked during the first 2 seconds of spout entry for rewarded and unrewarded trials during training and reversal, respectively. Decoders were constructed separately for each animal and used only neurons from that animal that were present on T2, T6 and R1 or on T2, T6 and R3. The decoder was trained on T2 and T6 activity. Activity of the population was Z-scored, then PCA was performed, keeping the top 20 principal components. before training LDA to predict reward status. Validation accuracy of the decoders was tested using leave-one-trial-out cross-validation. The decoder was then applied to either R1 or R3 activity, throughout 25 seconds of activity centered around spout entry. Prediction probabilities were used as a measure of confidence. High confidence in a prediction, either reward or no reward, was determined to be above 0.99 or below 0.01, respectively. Over the entire 25 seconds, contiguous periods of high confidence were identified. The number of switches between confidently predicting reward to confidently predicting no reward (and vice versa) were counted.

### Statistical procedures

All statistical tests were two sided and non-parametric unless stated otherwise. The Holm-Bonferroni method was used to adjust for multiple comparisons.

---

**Figure S1:**
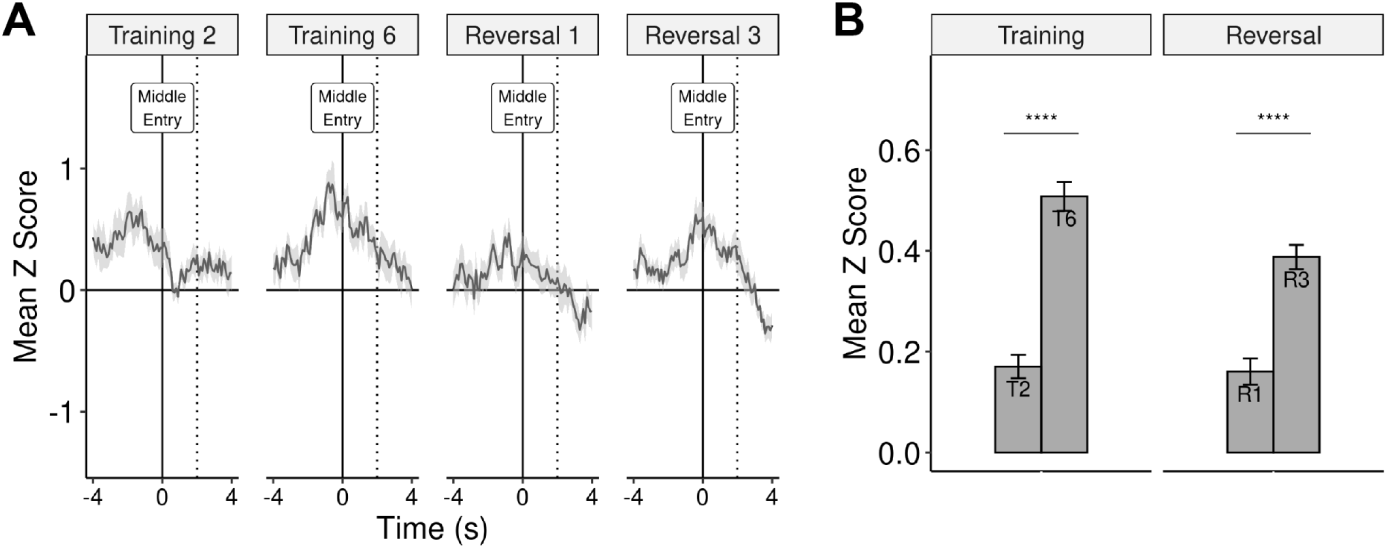
Additional LC fiber photometry recordings during the task. (A) Middle-entry aligned LC activity expressed as a *z* score (mean ± s.e.m.) across days. (B) Average LC activity (mean *z* score) during the first two seconds following middle entry (mean ± s.e.m.). Wilcoxon signed rank test on paired samples, ****p ≤ 0.0001. *n* = 6 mice.

**Figure S2:**
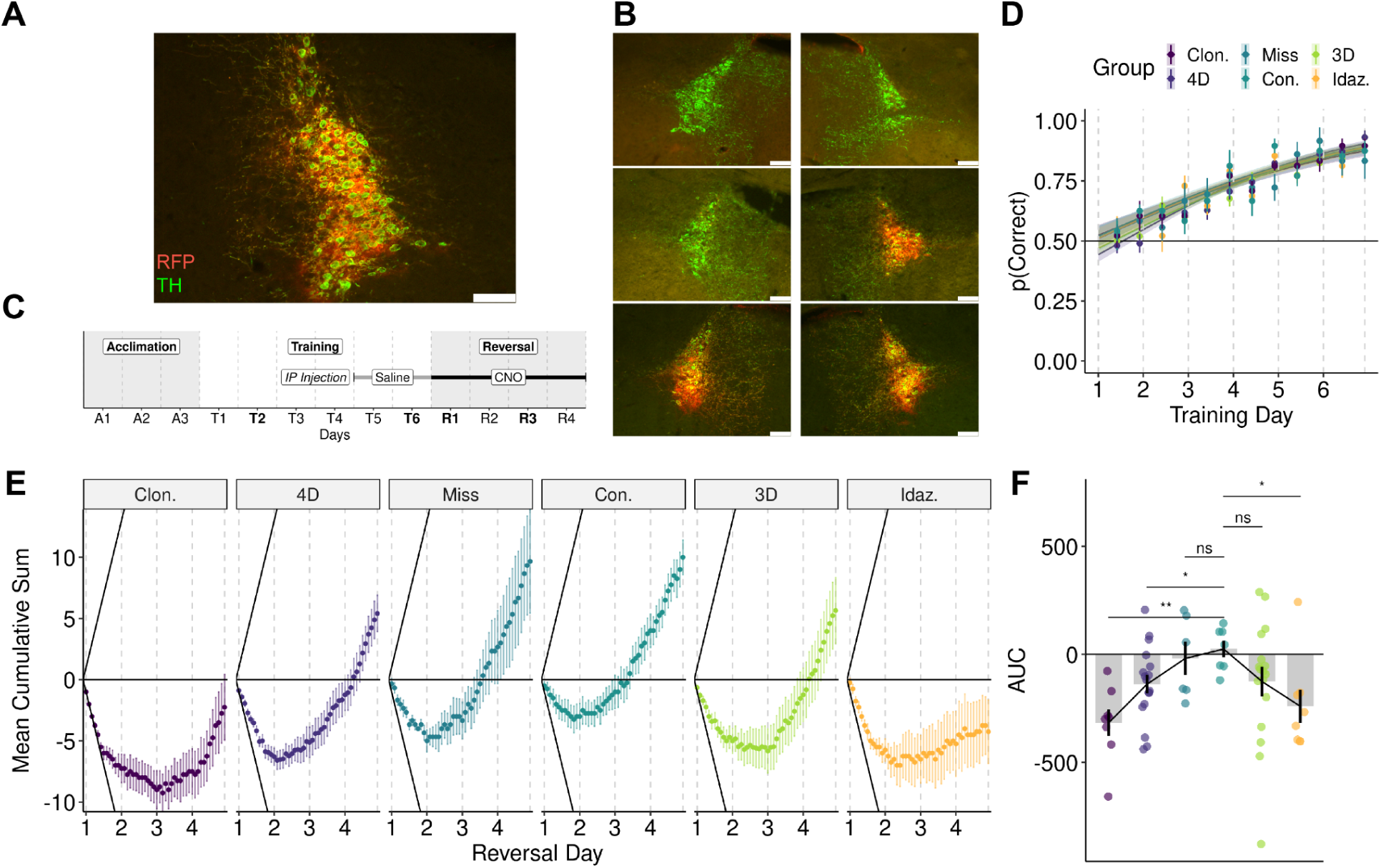
Chemogenetic manipulation of LC activity during reversal. (A) Representative image of DREADD-infected *Dbh*-expressing neurons (red) present within the LC (green). (B) Examples of varying DREADD expression (red) in the LC (green): miss (top), unilateral (middle), bilateral (bottom). Each row shows both LCs from a representative animal. (C) Experimental timeline schematic: Saline was injected (ip) on the final days of training. CNO was injected on each reversal day to activate the DREADD-expressing cells. (D) Task performance of all treatment groups during training (mean + s.e.m.): the proportion of rewarded trials/session (dots) and the proportion predicted with a logistic regression model (lines). Type 1 ANOVA chi-squared test, trial number: X^2^ (1) = 341.83, p < 0.0001, treatment group: X^2^ (5) = 1.79, p = 0.88, trial number x treatment group interaction: X^2^ (5) = 2.99, p = 0.70. (E) To further visualize behavior, trials were coded based on reward condition (+1 rewarded, -1 unrewarded) and the cumulative sum across all reversal trials was calculated for each animal and averaged across the groups (mean ± s.e.m.). For comparison, diagonal black lines represent the slope of the hypothetical mean cumulative sum if all trials were rewarded (upper) or unrewarded (lower). (F) The area under the cumulative sum curve (in E) for each animal (points) and the mean for each treatment group (bar + s.e.m.). Wilcoxon rank sum test with control as reference, *p ≤ 0.05, **p ≤ 0.01. *n* = 7 control, 8 clonidine, 8 idazoxan, 6 misses, 17 3D DREADD, and 17 4D DREADD mice.

**Figure S3:**
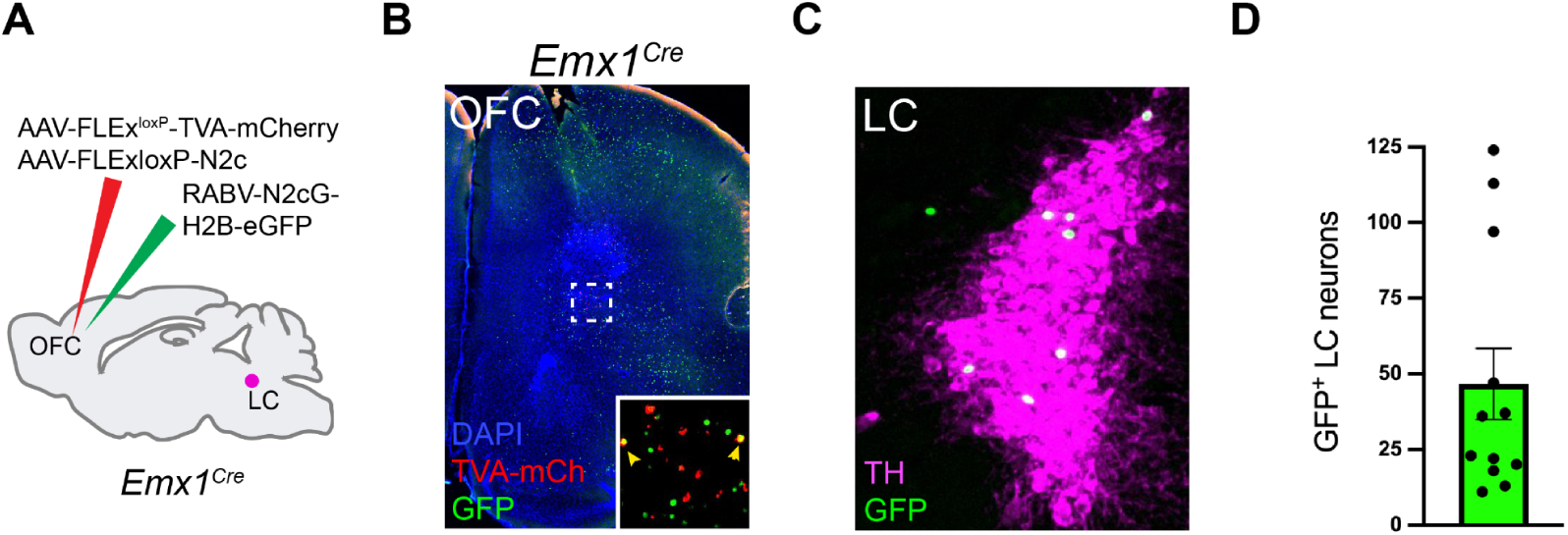
Monosynaptic rabies tracing identifies a direct projection from LC onto *Emx1^Cre^*^+^ OFC neurons. (A) Schematic for trans-synaptic rabies tracing from *Emx1^+^* OFC neurons. (B) Cre-dependent rabies helper AAVs expressing TVA receptor (red) and N2c glycoprotein were unilaterally co-injected into the OFC of *Emx1^Cre^* mice three weeks prior to injection of RABV-CVS-N2c^ΔG^-H2B-eGFP (green). Cell nuclei are counterstained with DAPI (blue). Yellow arrowheads point to putative OFC starter cells (inset). (C) Representative image of rabies-labeled input neurons (green) present within the LC (magenta) from an animal with *Emx1^Cre^* OFC starter cells. (D) Quantification of GFP labeled cells in the LC from *Emx1^Cre^*OFC tracing experiments (mean ± s.e.m). Points represent the sum cell count per animal from 5 50μm LC brain sections (approximately one third of the LC). *n* = 12 mice.

**Figure S4:**
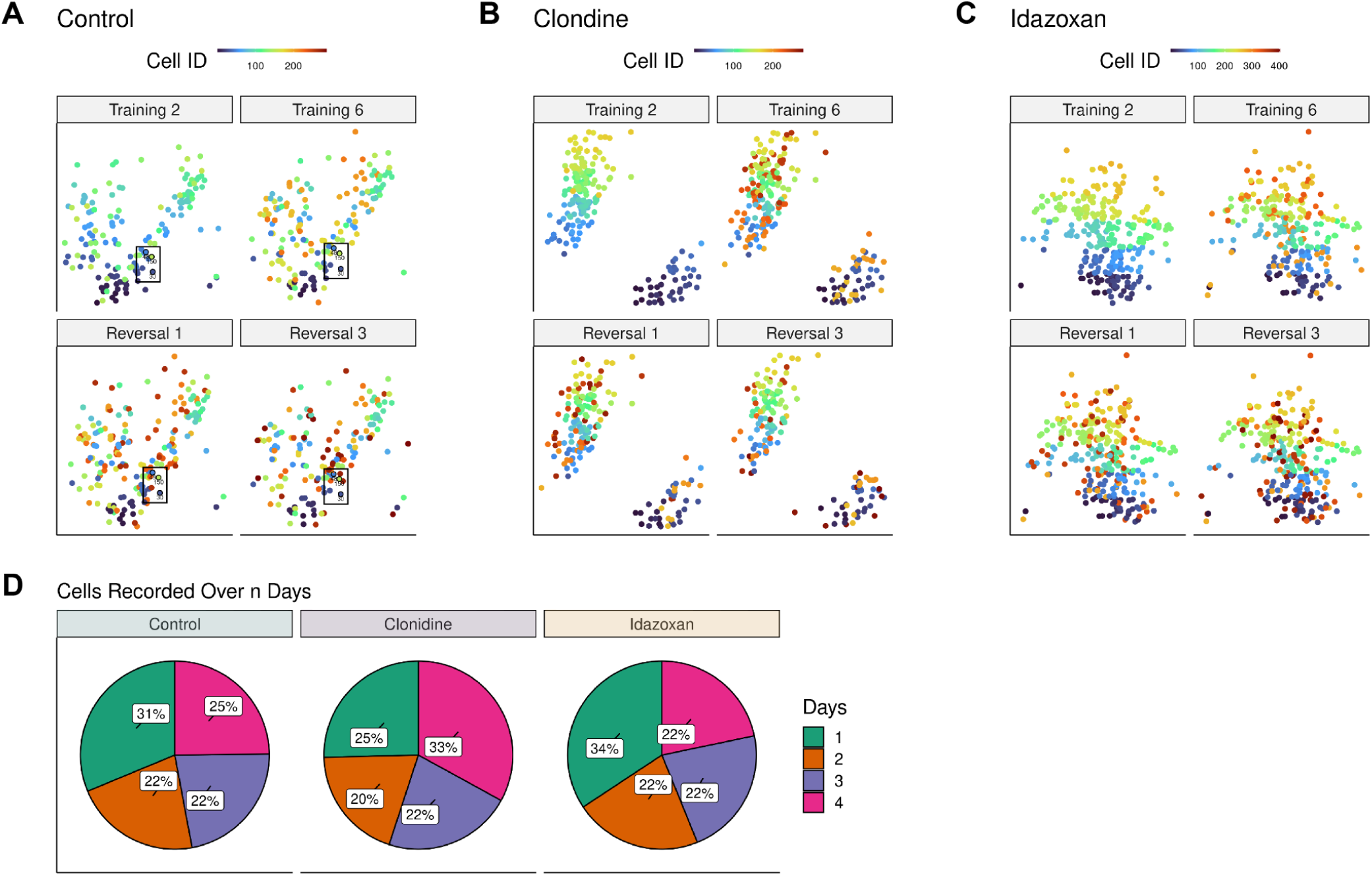
Comparison of the spatial location and presence of OFC cells across days. (A) Spatial location of OFC cells from an example control animal colored by cell identity. The color of each cell represents its unique, assigned ID number within each animal. As illustrated in the insets, cells recorded on more than one day have the same color in each panel. (B) Spatial location of OFC cells from an example clonidine animal colored by cell identity. (C) Spatial location of OFC cells from an example idazoxan animal colored by cell identity. (D) The distribution of the number of days that individual cells were recorded differed across treatment groups. Pearson’s chi-squared test, X^2^ (6) = 69.99, p < 0.0001. *n* = 1,519 cells (3 control mice), 1,694 cells (3 clonidine mice), 1,787 cells (3 idazoxan mice).

**Figure S5:**
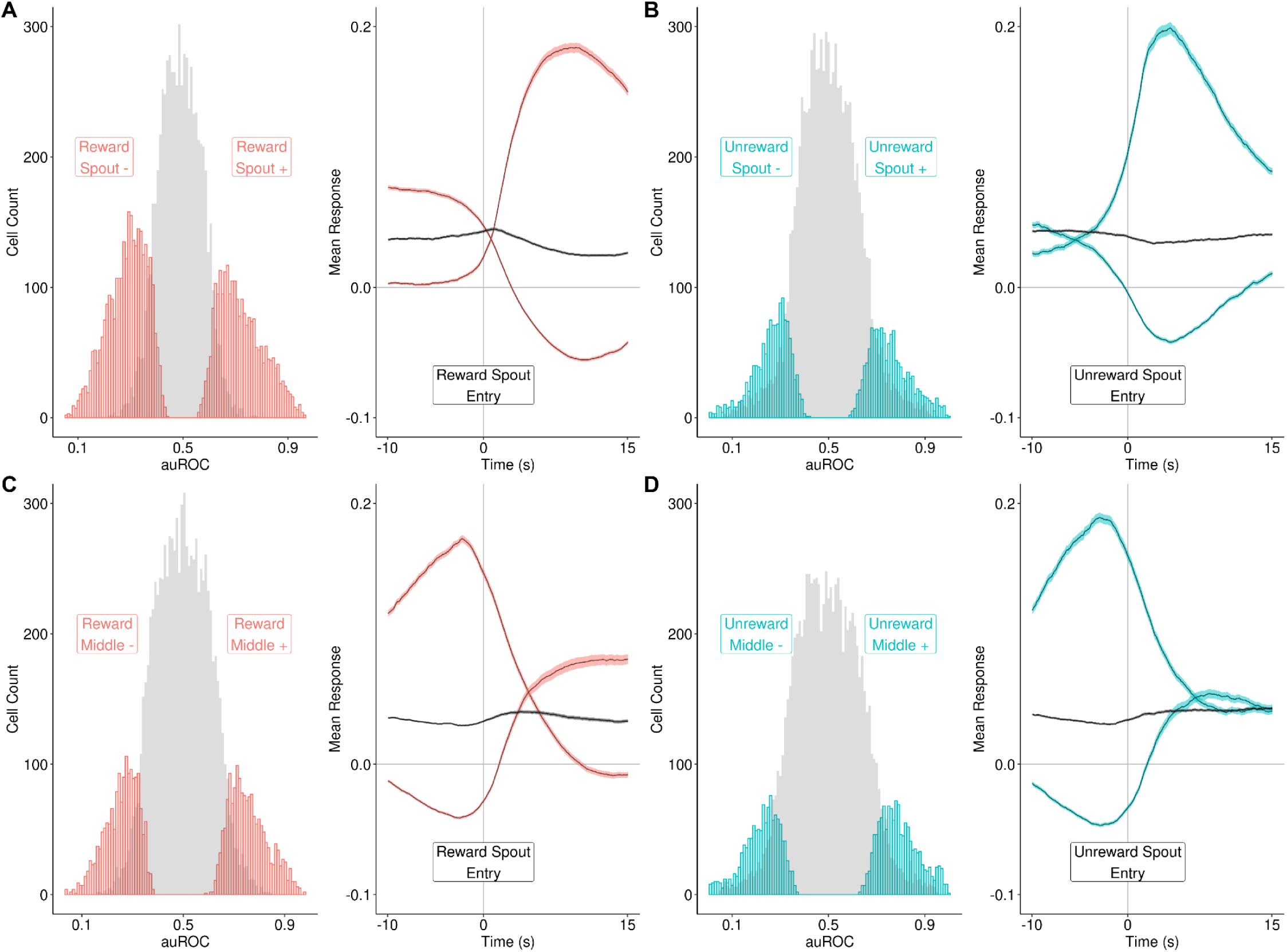
ROC-AUC Analysis. In **A-D**, each set of plots shows a histogram of the area under the receiver operator curve for all cellular responses to each region of interest (left) and the trial-averaged responses (mean ± s.e.m.) aligned to ROI-entry (right). Significant responses to the ROI (Methods) are indicated in color based on reward condition. *n* = 12,237 responses from 5,000 cells. **(A)** Rewarded Spout +, *n* = 2438 responses (19.9%). Rewarded Spout -, *n* = 3068 (25.1%). **(B)** Unrewarded Spout +, *n* = 1315 responses (10.7%), Unrewarded Spout -, *n* = 1363 responses (11.1%). **(C)** Rewarded Middle +, *n* = 1662 responses (13.6%). Rewarded Middle -, *n* = 1640 (13.4%). **(D)** Unrewarded Middle +, *n* = 1298 responses (10.6%), Unrewarded Middle -, *n* = 1185 responses (9.7%).

**Figure S6:**
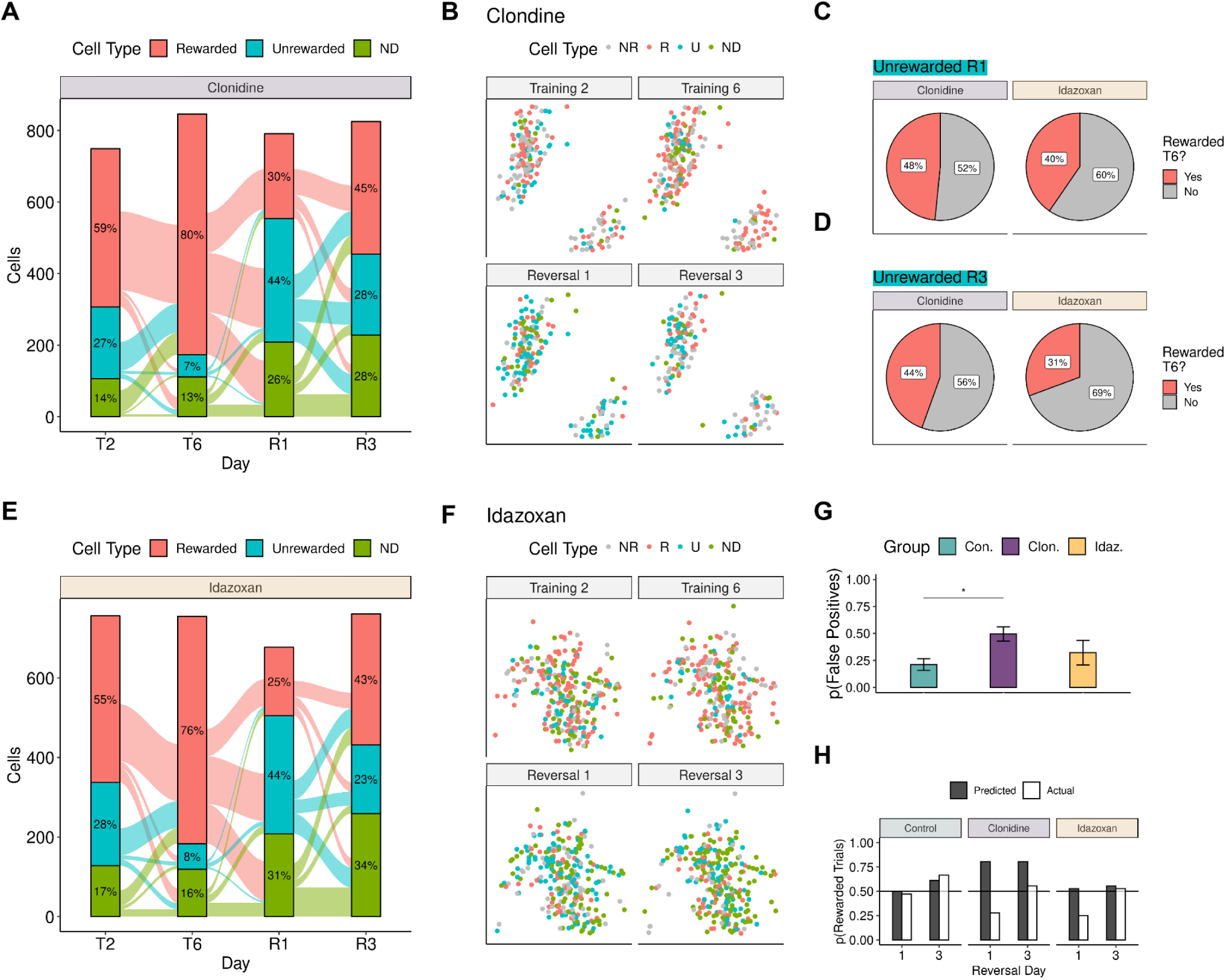
Effects of bidirectional alteration of LC activity on OFC task representation during reversal. (A) Alluvial plot showing how the distribution of cell types changes during training and reversal in the clonidine group. (B) Spatial location of OFC cells from an example clonidine animal colored by cell type. In C-D, plots compare the proportion of reversal *Unrewarded* cells that were part of the T6 *Rewarded* ensemble between the clonidine and idazoxan groups. (C) R1 comparison. Pearson’s chi-squared test, X^2^ (1) = 4.13, p = 0.04. (D) R3 comparison. Pearson’s chi-squared test, X^2^ (1) = 7.96, p = 0.005. (E) Alluvial plot showing how the distribution of cell types changes during training and reversal in the idazoxan group. (F) Spatial location of OFC cells from an example idazoxan animal colored by cell type. (G) The proportion of unrewarded reversal trials that were predicted as rewarded by the decoders. The proportion differed among the treatment groups. Kruskal-Wallis rank sum test, p=0.04. Wilcoxon rank sum test with control as reference, Clonidine: *p ≤ 0.05, Idazoxan: p ≥ 0.05. (H) Comparison across treatment groups of the proportion of total reversal trials that were predicted as rewarded by the decoders versus the proportion of total trials that were actually rewarded. *n* = 3 mice/group.

## References

1. Rich, E.L., and Wallis, J.D. (2016). Decoding subjective decisions from orbitofrontal cortex. Nat. Neurosci. 19, 973–980.

2. Riceberg, J.S., and Shapiro, M.L. (2017). Orbitofrontal Cortex Signals Expected Outcomes with Predictive Codes When Stable Contingencies Promote the Integration of Reward History. J. Neurosci. 37, 2010– 2021.

3. Ballesta, S., Shi, W., and Padoa-Schioppa, C. (2022). Orbitofrontal cortex contributes to the comparison of values underlying economic choices. Nat. Commun. 13, 4405.

4. Namboodiri, V.M.K., Otis, J.M., Heeswijk, K., Voets, E.S., Alghorazi, R.A., Rodriguez-Romaguera, J., Mihalas, S., and Stuber, G.D. (2019). Single-cell activity tracking reveals that orbitofrontal neurons acquire and maintain a long-term memory to guide behavioral adaptation. Nat. Neurosci., 1–19.

5. Basu, R., Gebauer, R., Herfurth, T., Kolb, S., Golipour, Z., Tchumatchenko, T., and Ito, H.T. (2021). The orbitofrontal cortex maps future navigational goals. Nature 599, 1–4.

6. Hirokawa, J., Vaughan, A., Masset, P., Ott, T., and Kepecs, A. (2019). Frontal cortex neuron types categorically encode single decision variables. Nature 148, 1–28.

7. Padoa-Schioppa, C., and Assad, J.A. (2006). Neurons in the orbitofrontal cortex encode economic value. Nature 441, 223–226.

8. Wilson, R.C., Takahashi, Y.K., Schoenbaum, G., and Niv, Y. (2014). Orbitofrontal cortex as a cognitive map of task space. Neuron 81, 267–279.

9. Rolls, E.T., Critchley, H.D., Mason, R., and Wakeman, E.A. (1996). Orbitofrontal cortex neurons: role in olfactory and visual association learning. J. Neurophysiol. 75, 1970–1981.

10. Gardner, M.P.H., and Schoenbaum, G. (2021). The orbitofrontal cartographer. Behav. Neurosci. 135, 267–276.

11. Knudsen, E.B., and Wallis, J.D. (2022). Taking stock of value in the orbitofrontal cortex. Nat. Rev. Neurosci. 23, 428–438.

12. Niv, Y. (2019). Learning task-state representations. Nat. Neurosci. 22, 1544–1553.

13. Rudebeck, P.H., and Murray, E.A. (2014). The orbitofrontal oracle: cortical mechanisms for the prediction and evaluation of specific behavioral outcomes. Neuron 84, 1143–1156.

14. Sadacca, B.F., Wikenheiser, A.M., and Schoenbaum, G. (2017). Toward a theoretical role for tonic norepinephrine in the orbitofrontal cortex in facilitating flexible learning. Neuroscience 345, 124–129.

15. Aston-Jones, G., and Cohen, J.D. (2005). An integrative theory of locus coeruleus-norepinephrine function: adaptive gain and optimal performance. Annu. Rev. Neurosci. 28, 403–450.

16. Bouret, S., and Sara, S.J. (2005). Network reset: a simplified overarching theory of locus coeruleus noradrenaline function. Trends Neurosci. 28, 574–582.

17. Yu, A.J., and Dayan, P. (2005). Uncertainty, neuromodulation, and attention. Neuron 46, 681–692.

18. Sales, A.C., Friston, K.J., Jones, M.W., Pickering, A.E., and Moran, R.J. (2019). Locus Coeruleus tracking of prediction errors optimises cognitive flexibility: An Active Inference model. PLoS Comput. Biol. 15, e1006267.

19. Gu, Q. (2002). Neuromodulatory transmitter systems in the cortex and their role in cortical plasticity. Neuroscience 111, 815–835.

20. Seu, E., Lang, A., Rivera, R.J., and Jentsch, J.D. (2009). Inhibition of the norepinephrine transporter improves behavioral flexibility in rats and monkeys. Psychopharmacology 202, 505–519.

21. Devauges, V., and Sara, S.J. (1990). Activation of the noradrenergic system facilitates an attentional shift in the rat. Behav. Brain Res. 39, 19–28.

22. Cope, Z.A., Vazey, E.M., Floresco, S.B., and Aston Jones, G.S. (2019). DREADD-mediated modulation of locus coeruleus inputs to mPFC improves strategy set-shifting. Neurobiol. Learn. Mem. 161, 1–11.

23. Aston-Jones, G., Rajkowski, J., and Cohen, J. (1999). Role of locus coeruleus in attention and behavioral flexibility. Biol. Psychiatry 46, 1309–1320.

24. Rorabaugh, J.M., Chalermpalanupap, T., Botz-Zapp, C.A., Fu, V.M., Lembeck, N.A., Cohen, R.M., and Weinshenker, D. (2017). Chemogenetic locus coeruleus activation restores reversal learning in a rat model of Alzheimer’s disease. Brain 140, 3023–3038.

25. McBurney-Lin, J., Vargova, G., Garad, M., Zagha, E., and Yang, H. (2022). The locus coeruleus mediates behavioral flexibility. Cell Rep. 41, 111534.

26. Breton-Provencher, V., Drummond, G.T., Feng, J., Li, Y., and Sur, M. (2022). Spatiotemporal dynamics of noradrenaline during learned behaviour. Nature 606, 732–738.

27. Tervo, D.G.R., Proskurin, M., Manakov, M., Kabra, M., Vollmer, A., Branson, K., and Karpova, A.Y. (2014). Behavioral Variability through Stochastic Choice and Its Gating by Anterior Cingulate Cortex. Cell 159, 21–32.

28. Jordan, R., and Keller, G.B. (2023). The locus coeruleus broadcasts prediction errors across the cortex to promote sensorimotor plasticity. Elife 12. 10.7554/eLife.85111.

29. Berridge, C.W., and Waterhouse, B.D. (2003). The locus coeruleus-noradrenergic system: modulation of behavioral state and state-dependent cognitive processes. Brain Res. Brain Res. Rev. 42, 33–84.

30. Torregrossa, M.M., Quinn, J.J., and Taylor, J.R. (2008). Impulsivity, compulsivity, and habit: the role of orbitofrontal cortex revisited. Biol. Psychiatry 63, 253–255.

31. Fettes, P., Schulze, L., and Downar, J. (2017). Cortico-Striatal-Thalamic Loop Circuits of the Orbitofrontal Cortex: Promising Therapeutic Targets in Psychiatric Illness. Front. Syst. Neurosci. 11, 25.

32. Hyman, S.E. (2005). Addiction: a disease of learning and memory. Am. J. Psychiatry 162, 1414–1422.

33. Uddin, L.Q. (2021). Cognitive and behavioural flexibility: neural mechanisms and clinical considerations. Nat. Rev. Neurosci. 22, 167–179.

34. Tillage, R.P., Sciolino, N.R., Plummer, N.W., Lustberg, D., Liles, L.C., Hsiang, M., Powell, J.M., Smith, K.G., Jensen, P., and Weinshenker, D. (2020). Elimination of galanin synthesis in noradrenergic neurons reduces galanin in select brain areas and promotes active coping behaviors. Brain Struct. Funct. 225, 785–803.

35. Aston-Jones, G., Rajkowski, J., and Kubiak, P. (1997). Conditioned responses of monkey locus coeruleus neurons anticipate acquisition of discriminative behavior in a vigilance task. Neuroscience 80, 697–715.

36. Xiang, L., Harel, A., Gao, H., Pickering, A.E., Sara, S.J., and Wiener, S.I. (2019). Behavioral correlates of activity of optogenetically identified locus coeruleus noradrenergic neurons in rats performing T-maze tasks. Sci. Rep. 9, 1361.

37. Su, Z., and Cohen, J.Y. (2022). Two types of locus coeruleus norepinephrine neurons drive reinforcement learning. bioRxiv, 2022.12.08.519670. 10.1101/2022.12.08.519670.

38. den Ouden, H.E.M., Kok, P., and de Lange, F.P. (2012). How prediction errors shape perception, attention, and motivation. Front. Psychol. 3, 548.

39. Sciolino, N.R., Hsiang, M., Mazzone, C.M., Wilson, L.R., Plummer, N.W., Amin, J., Smith, K.G., McGee, C.A., Fry, S.A., Yang, C.X., et al. (2022). Natural locus coeruleus dynamics during feeding. Sci Adv 8, eabn9134.

40. McBurney-Lin, J., Sun, Y., Tortorelli, L.S., Nguyen, Q.A.T., Haga-Yamanaka, S., and Yang, H. (2020). Bidirectional pharmacological perturbations of the noradrenergic system differentially affect tactile detection. Neuropharmacology 174, 108151.

41. Simson, P.E., and Weiss, J.M. (1987). Alpha-2 receptor blockade increases responsiveness of locus coeruleus neurons to excitatory stimulation. J. Neurosci. 7, 1732–1740.

42. Yerkes, R.M., and Dodson, J.D. (1908). The relation of strength of stimulus to rapidity of habit-formation. J. Comp. Neurol. Psychol. 18, 459–482.

43. Elorette, C., Fujimoto, A., Fredericks, J.M., Stoll, F.M., Russ, B.E., and Rudebeck, P.H. (2021). Piecing together the orbitofrontal puzzle. Behav. Neurosci. 135, 301–311.

44. Stalnaker, T.A., Cooch, N.K., and Schoenbaum, G. (5/2015). What the orbitofrontal cortex does not do. Nat. Neurosci. 18, 620–627.

45. Zhou, J., Jia, C., Montesinos-Cartagena, M., Gardner, M.P.H., Zong, W., and Schoenbaum, G. (2021). Evolving schema representations in orbitofrontal ensembles during learning. Nature 590, 606–611.

46. Lopatina, N., Sadacca, B.F., McDannald, M.A., Styer, C.V., Peterson, J.F., Cheer, J.F., and Schoenbaum, G. (2017). Ensembles in medial and lateral orbitofrontal cortex construct cognitive maps emphasizing different features of the behavioral landscape. Behav. Neurosci. 131, 201–212.

47. Stalnaker, T.A., Raheja, N., and Schoenbaum, G. (2021). Orbitofrontal State Representations Are Related to Choice Adaptations and Reward Predictions. J. Neurosci. 41, 1941–1951.

48. Sul, J.H., Kim, H., Huh, N., Lee, D., and Jung, M.W. (05/2010). Distinct Roles of Rodent Orbitofrontal and Medial Prefrontal Cortex in Decision Making. Neuron 66, 449–460.

49. Thorpe, S.J., Rolls, E.T., and Maddison, S. (1/1983). The orbitofrontal cortex: Neuronal activity in the behaving monkey. Exp. Brain Res. 49. 10.1007/BF00235545.

50. Takahashi, Y.K., Roesch, M.R., Stalnaker, T.A., Haney, R.Z., Calu, D.J., Taylor, A.R., Burke, K.A., and Schoenbaum, G. (04/2009). The Orbitofrontal Cortex and Ventral Tegmental Area Are Necessary for Learning from Unexpected Outcomes. Neuron 62, 269–280.

51. Matias, S., Lottem, E., Dugué, G.P., and Mainen, Z.F. (2017). Activity patterns of serotonin neurons underlying cognitive flexibility. Elife 6. 10.7554/eLife.20552.

52. Lerner, T.N., Holloway, A.L., and Seiler, J.L. (2021). Dopamine, Updated: Reward Prediction Error and Beyond. Curr. Opin. Neurobiol. 67, 123–130.

53. Grossman, C.D., Bari, B.A., and Cohen, J.Y. (2022). Serotonin neurons modulate learning rate through uncertainty. Curr. Biol. 32, 586–599.e7.

54. Hollerman, J.R., and Schultz, W. (1998). Dopamine neurons report an error in the temporal prediction of reward during learning. Nat. Neurosci. 1, 304–309.

55. Radke, A.K., Kocharian, A., Covey, D.P., Lovinger, D.M., Cheer, J.F., Mateo, Y., and Holmes, A. (2019). Contributions of nucleus accumbens dopamine to cognitive flexibility. Eur. J. Neurosci. 50, 2023–2035.

56. Clarke, H.F., Dalley, J.W., Crofts, H.S., Robbins, T.W., and Roberts, A.C. (2004). Cognitive inflexibility after prefrontal serotonin depletion. Science 304, 878–880.

57. Crofts, H.S., Dalley, J.W., Collins, P., Van Denderen, J.C., Everitt, B.J., Robbins, T.W., and Roberts, A.C. (2001). Differential effects of 6-OHDA lesions of the frontal cortex and caudate nucleus on the ability to acquire an attentional set. Cereb. Cortex 11, 1015–1026.

58. Boulougouris, V., Glennon, J.C., and Robbins, T.W. (2008). Dissociable effects of selective 5-HT2A and 5-HT2C receptor antagonists on serial spatial reversal learning in rats. Neuropsychopharmacology 33, 2007–2019.

59. Furr, A., Lapiz-Bluhm, M.D., and Morilak, D.A. (2012). 5-HT2A receptors in the orbitofrontal cortex facilitate reversal learning and contribute to the beneficial cognitive effects of chronic citalopram treatment in rats. Int. J. Neuropsychopharmacol. 15, 1295–1305.

60. Arnsten, A.F., and Goldman-Rakic, P.S. (1985). Alpha 2-adrenergic mechanisms in prefrontal cortex associated with cognitive decline in aged nonhuman primates. Science 230, 1273–1276.

61. Zerbi, V., Floriou-Servou, A., Markicevic, M., Vermeiren, Y., Sturman, O., Privitera, M., von Ziegler, L., Ferrari, K.D., Weber, B., De Deyn, P.P., et al. (2019). Rapid Reconfiguration of the Functional Connectome after Chemogenetic Locus Coeruleus Activation. Neuron 103, 702–718.e5.

62. Glennon, E., Valtcheva, S., Zhu, A., Wadghiri, Y.Z., Svirsky, M.A., and Froemke, R.C. (2023). Locus coeruleus activity improves cochlear implant performance. Nature 613, 317–323.

63. Perlman, K., Benrimoh, D., Israel, S., Rollins, C., Brown, E., Tunteng, J.-F., You, R., You, E., Tanguay-Sela, M., Snook, E., et al. (2019). A systematic meta-review of predictors of antidepressant treatment outcome in major depressive disorder. J. Affect. Disord. 243, 503–515.

64. Rush, A.J., Trivedi, M.H., Wisniewski, S.R., Stewart, J.W., Nierenberg, A.A., Thase, M.E., Ritz, L., Biggs, M.M., Warden, D., Luther, J.F., et al. (2006). Bupropion-SR, sertraline, or venlafaxine-XR after failure of SSRIs for depression. N. Engl. J. Med. 354, 1231–1242.

65. Havekes, R., Nijholt, I.M., Luiten, P.G.M., and Van der Zee, E.A. (2006). Differential involvement of hippocampal calcineurin during learning and reversal learning in a Y-maze task. Learn. Mem. 13, 753– 759.

66. Bogovic, J.A., Hanslovsky, P., Wong, A., and Saalfeld, S. (2016). Robust registration of calcium images by learned contrast synthesis. In 2016 IEEE 13th International Symposium on Biomedical Imaging (ISBI), pp. 1123–1126.

67. Lerner, T.N., Shilyansky, C., Davidson, T.J., Evans, K.E., Beier, K.T., Zalocusky, K.A., Crow, A.K., Malenka, R.C., Luo, L., Tomer, R., et al. (2015). Intact-Brain Analyses Reveal Distinct Information Carried by SNc Dopamine Subcircuits. Cell 162, 635–647.

68. Li, L., Feng, X., Zhou, Z., Zhang, H., Shi, Q., Lei, Z., Shen, P., Yang, Q., Zhao, B., Chen, S., et al. (2018). Stress Accelerates Defensive Responses to Looming in Mice and Involves a Locus Coeruleus-Superior Colliculus Projection. Curr. Biol. 28, 859–871.e5.

69. Giovannucci, A., Friedrich, J., Gunn, P., Kalfon, J., Brown, B.L., Koay, S.A., Taxidis, J., Najafi, F., Gauthier, J.L., Zhou, P., et al. (2019). CaImAn an open source tool for scalable calcium imaging data analysis. Elife 8. 10.7554/eLife.38173.

70. Kingsbury, L., Huang, S., Raam, T., Ye, L.S., Wei, D., Hu, R.K., Ye, L., and Hong, W. (2020). Cortical Representations of Conspecific Sex Shape Social Behavior. Neuron 107, 941–953.e7.

71. Timme, N.M., and Lapish, C. (2018). A Tutorial for Information Theory in Neuroscience. eNeuro 5. 10.1523/ENEURO.0052-18.2018.

